# Early intermediates in bacterial RNA polymerase promoter melting visualized by time-resolved cryo-electron microscopy

**DOI:** 10.1101/2024.03.13.584744

**Authors:** Ruth M. Saecker, Andreas U. Mueller, Brandon Malone, James Chen, William C. Budell, Venkata P. Dandey, Kashyap Maruthi, Joshua H. Mendez, Nina Molina, Edward T. Eng, Laura Y. Yen, Clinton S. Potter, Bridget Carragher, Seth A. Darst

## Abstract

During formation of the transcription-competent open complex (RPo) by bacterial RNA polymerases (RNAP), transient intermediates pile up before overcoming a rate-limiting step. Structural descriptions of these interconversions in real time are unavailable. To address this gap, time-resolved cryo-electron microscopy (cryo-EM) was used to capture four intermediates populated 120 or 500 milliseconds (ms) after mixing *Escherichia coli* σ^70^-RNAP and the αP_R_ promoter. Cryo-EM snapshots revealed the upstream edge of the transcription bubble unpairs rapidly, followed by stepwise insertion of two conserved nontemplate strand (nt-strand) bases into RNAP pockets. As nt-strand “read-out” extends, the RNAP clamp closes, expelling an inhibitory σ^70^ domain from the active-site cleft. The template strand is fully unpaired by 120 ms but remains dynamic, indicating yet unknown conformational changes load it in subsequent steps. Because these events likely describe DNA opening at many bacterial promoters, this study provides needed insights into how DNA sequence regulates steps of RPo formation.

---

Initiation of transcription is a major control point for regulating gene expression in all cells. In bacteria, a single catalytic core RNAP (E, subunit composition α_2_ββ’ω) performs all transcription but must combine with a σ factor, such as σ^70^ in *Escherichia coli* (*Eco*), to form the holoenzyme (Eσ^70^) for initiation ^1,2^. The RNAP structure resembles a crab claw with pincers formed from the large β and β’ subunits ^3^. The enzyme active site, marked by a bound Mg^2+^-ion, lies deep in the large cleft between the pincers.

In initiation, interactions between Eσ^70^ and specific promoter DNA sequences (most importantly the -35 and -10 elements ^4^, see Extended Data Fig. 1a for the λP_R_ promoter sequence used here) trigger a series of isomerization steps that separate the DNA strands within the ∼13 nucleotide transcription bubble ^5,6^. Ultimately, a transcription-competent “open” promoter complex (RPo) forms in which the DNA template-strand (t-strand) is positioned deep (∼70 Å) within the RNAP active-site cleft with bases at +1 (start site) and +2 positioned to template incoming NTP substrates ^7^. Correct alignment of the first two NTPs with respect to the active site Mg^2+^-ions catalyzes phosphodiester bond formation ^8^. Thus, RPo formation is required, and is often the rate-limiting step, for the initiation of every RNA chain ^9^.

Promoter opening is a multi-step process during which transient intermediates appear and disappear during progression to the final RPo ^5,6,10,11^. Initial recognition of the promoter -35 element by σ^70^ domain 4 (σ^70^)^12^ outside the RNAP cleft positions the promoter -10 element near the upstream entrance to the cleft. ‘Nucleation’ of the transcription bubble is thought to occur by a ‘flip-and-capture’ mechanism, whereby bases of the -10 element nontemplate-strand [nt-strand; namely A_-11_(nt) and T_-7_(nt), Extended Data Fig. 1a] flip-out of the duplex DNA base stack and are captured in cognate pockets of σ^70^ ^13^. The transcription bubble then propagates in the downstream direction to encompass the transcription start site (+1). Formation of the final RPo involves: 1) Isomerization of an invariant W-dyad of σ^70^ (W433/W434) from an edge-on to a chair-like conformation, which stabilizes the upstream edge of the bubble, and 2) loading of the resulting single-stranded DNA and downstream duplex DNA into the RNAP cleft ^10,14^.

In *Eco*, promoter DNA loading in the RNAP cleft requires the expulsion of the N-terminal domain 1.1 of σ^70^, σ^70^ ^15,16^. Conserved in many bacterial group 1 σ’s^17^, σ^70^ folds into a negatively-charged four-helix bundle that protects the highly basic RNAP cleft from inappropriately interacting with non-specific nucleic acid. Ejection of σ^70^ serves as a critical regulatory step in RPo formation, but when and how σ^70^ is displaced by incoming promoter DNA remains unknown.

Decades of biochemical and biophysical studies have provided mechanistic models for RPo formation, and Eσ^70^ and RPo structures have provided constraints on those models ^5,6,18^. Advances in cryo-electron microscopy (cryo-EM) that allow high-resolution information to be extracted from dynamic, heterogeneous samples enabled the first inroads into structural analysis of intermediates in RPo formation ^10,19^. Use of standard cryo-EM sample preparation methods in these studies necessitated conditions (e.g., destabilizing factors, promoter mutants) where intermediates were populated at equilibrium. However, formation of RPo is intrinsically a nonequilibrium process. Observing unperturbed RPo intermediates requires high-resolution visualization of RPo formation in real time. To achieve this, we took advantage of two advances in cryo-EM methodology: i) The development of cryo-EM instrumentation that allows mixing and capture of biomolecules on a subsecond time scale (time-resolved-Spotiton, or tr-Spotiton; Extended Data Fig. 1b) ^20^; and ii) Computational approaches that resolve the conformational heterogeneity in single particle images (3D-variability analysis, 3DVA) ^21^. The results of detailed mechanistic studies of RPo formation at the σP_R_ promoter ^5,6^ allowed the choice of times and conditions where early intermediates are predicted to pile up before the rate-limiting step (Fig. 1a). Together these advances allowed us to capture high-resolution structures of RPo intermediates as they formed in real time.

**Fig. 1.**
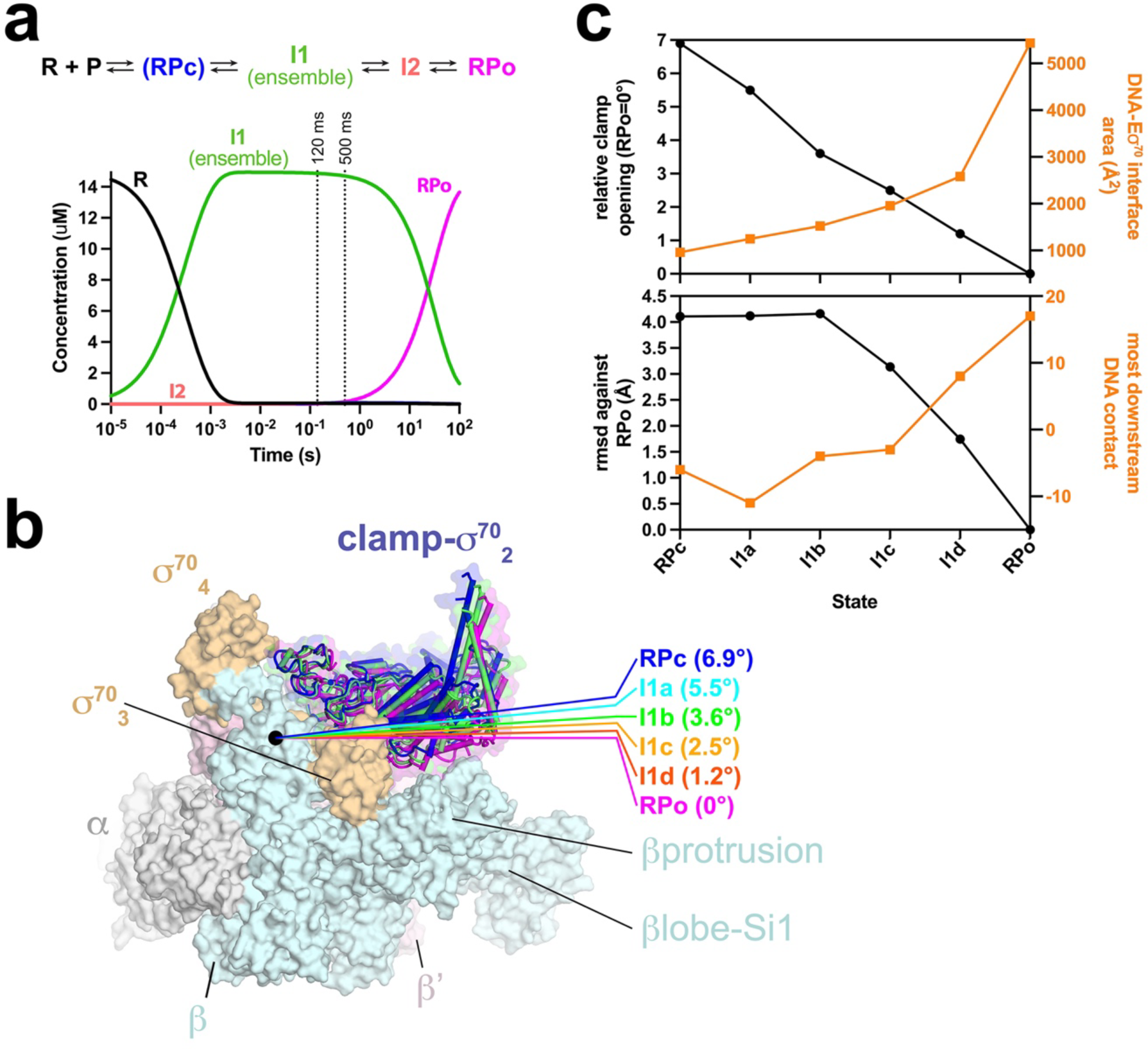
Promoter melting Intermediates on the σP_R_ Promoter. **a.** (*top*) Minimal kinetic mechanism for the formation of RPo on σP_R_, using the nomenclature of Record & colleagues ^5,6^, with two kinetically significant intermediates, I1 and I2. In this scheme RNAP (R) binds the σP_R_ promoter (P) and forms I1, an ensemble of rapidly equilibrating states ^5^. I1 converts to I2 in a rate-limiting step, which then converts rapidly to RPo. The intermediate RPc is only observed < 7°C. (*bottom*) Simulation of the time-course of the reaction under the conditions of the tr-Spotiton experiments (RT, [Eσ^70^] = 15 μM, [σP_R_-DNA] = 30 μM). See Methods for the kinetic parameters used to generate the simulation ^24^. For both 120 and 500 ms mixing times, only I1 (with no RPo) was expected. **b.** RNAP clamp conformational changes for intermediates determined in this work. The RPo structure (7MKD) ^7^ was used as a reference to superimpose the intermediate structures via σ-carbon atoms of the RNAP structural core, revealing a common RNAP structure (shown as a molecular surface) but with clamp conformational changes characterized as rigid body rotations about a rotation axis perpendicular to the page (denoted by the black dot). The clamp modules for RPc, I1b, and I1d are shown as backbone cartoons with cylindrical helices (σ^70^_NCR_ is omitted for clarity). The angles of clamp opening for all the intermediates are shown relative to RPo (0°). **c.** Structural properties used to order the complexes in the RPo formation pathway. (*top panel*) Plotted in black (left scale) is the clamp opening angle [relative to σP_R_-RPo (7MKD) ^7^ defined as 0°]. Plotted in orange (right scale) is the DNA-Eσ^70^ interface area (Å^2^) ^74^. (*bottom panel*) Plotted in black (left scale) is the root-mean-square deviation of σ-carbon positions (Å) for each complex superimposed with RPo. Plotted in orange (right scale) is the most downstream Eσ^70^-DNA contact observed in each complex. Also see Extended Data Fig. 1 and Supplementary Video 1.

## Results

### Direct visualization of DNA melting intermediates by tr-Spotiton

In the tr-Spotiton robot design, piezo dispensing tips direct two separate streams of ∼50 pL droplets onto a nanowire (’self-wicking’) cryo-EM grid ^22^ as it traverses towards vitrification, resulting in a stripe of sample across each grid (Extended Data Fig. 1b) ^20^. Complete mixing of the two samples occurs within ∼10 ms of colliding with the grid surface ^23^. Reaction times before vitrification can be varied by changing the velocity of the grid (Supplementary Video 1).

To trap RPo formation intermediates, *Eco* Eσ^70^ (∼30 μM) and a σP_R_ promoter DNA fragment (60 μM; Extended Data Fig. 1a) were deposited onto a self-wicking cryo-EM grid at room temperature (RT) from separate piezo tips (Extended Data Fig. 1b). On-grid mixing occurred at RT under buffer conditions where the kinetics of RPo formation have been well-characterized (Fig. 1a) ^5,24,25^ except 8 mM CHAPSO was present (in both samples) to eliminate particle orientation bias at the liquid-air interface ^26^. Under these conditions, the rate of RPo formation is expected to be largely determined by the rate-limiting step (I1 -> I2, k_2_ ∼0.04 s^-1^; Fig. 1a) ^25^. *In vitro* mechanistic studies predict an ensemble of early intermediates in rapid equilibrium with each other (I1 ensemble) between about 1 ms to 1 s after mixing (Fig. 1a) ^5,25,27^, while relaxation to RPo takes tens of seconds (the conversion from I2 to RPo is extremely fast so I2 is not populated under these conditions; Fig. 1a).

Multiple 120 ms mixing experiments and subsequent cryo-EM data collections were conducted. Of these, three datasets (datasets 1_120ms_, 2_120ms_, and 3_120ms_) were of high quality, yielding consensus structures with nominal resolutions from 3.3 to 3.4 Å (Table 1; Extended Data Fig. 2). A fourth dataset with a longer mixing time was also collected (dataset 4_500ms_) and processed, yielding a consensus structure with a nominal resolution of 3.0 Å (Table 1, Extended Data Fig. 3).

**Table 1.**
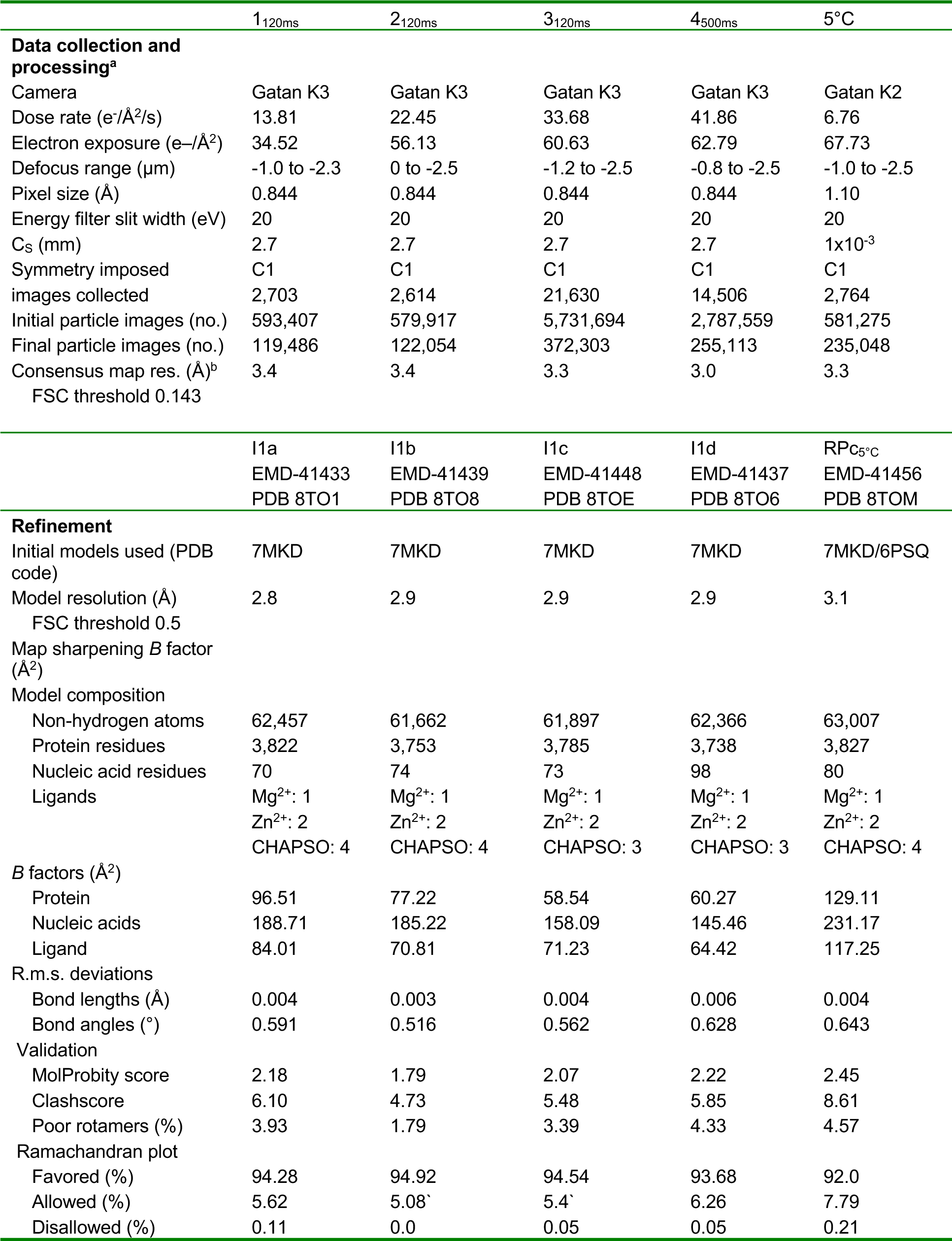
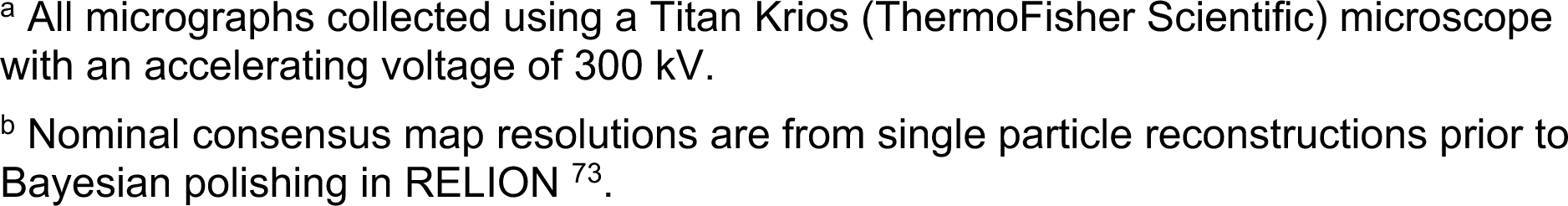
Cryo-EM data collection, refinement, and validation statistics.

|  | 1<sub>120ms</sub> | 2<sub>120ms</sub> | 3<sub>120ms</sub> | 4<sub>500ms</sub> | 5°C |
| --- | --- | --- | --- | --- | --- |
| **Data collection and processing<sup>a</sup>** |  |  |  |  |  |
| Camera | Gatan K3 | Gatan K3 | Gatan K3 | Gatan K3 | Gatan K2 |
| Dose rate (e<sup>-</sup>/Å<sup>2</sup>/s) | 13.81 | 22.45 | 33.68 | 41.86 | 6.76 |
| Electron exposure (e<sup>-</sup>/Å<sup>2</sup>) | 34.52 | 56.13 | 60.63 | 62.79 | 67.73 |
| Defocus range (μm) | -1.0 to -2.3 | 0 to -2.5 | -1.2 to -2.5 | -0.8 to -2.5 | -1.0 to -2.5 |
| Pixel size (Å) | 0.844 | 0.844 | 0.844 | 0.844 | 1.10 |
| Energy filter slit width (eV) | 20 | 20 | 20 | 20 | 20 |
| Cs (mm) | 2.7 | 2.7 | 2.7 | 2.7 | 1x10<sup>-3</sup> |
| Symmetry imposed | C1 | C1 | C1 | C1 | C1 |
| images collected | 2,703 | 2,614 | 21,630 | 14,506 | 2,764 |
| Initial particle images (no.) | 593,407 | 579,917 | 5,731,694 | 2,787,559 | 581,275 |
| Final particle images (no.) | 119,486 | 122,054 | 372,303 | 255,113 | 235,048 |
| Consensus map res. (Å)<sup>b</sup> | 3.4 | 3.4 | 3.3 | 3.0 | 3.3 |
| FSC threshold 0.143 |  |  |  |  |  |
|  | I1a | I1b | I1c | I1d | RPC<sub>5°C</sub> |
|  | EMD-41433 | EMD-41439 | EMD-41448 | EMD-41437 | EMD-41456 |
|  | PDB 8TO1 | PDB 8TO8 | PDB 8TOE | PDB 8TO6 | PDB 8TOM |
| **Refinement** |  |  |  |  |  |
| Initial models used (PDB code) | 7MKD | 7MKD | 7MKD | 7MKD | 7MKD/6PSQ |
| Model resolution (Å) | 2.8 | 2.9 | 2.9 | 2.9 | 3.1 |
| FSC threshold 0.5 |  |  |  |  |  |
| Map sharpening *B* factor (Å<sup>2</sup>) |  |  |  |  |  |
| Model composition |  |  |  |  |  |
| Non-hydrogen atoms | 62,457 | 61,662 | 61,897 | 62,366 | 63,007 |
| Protein residues | 3,822 | 3,753 | 3,785 | 3,738 | 3,827 |
| Nucleic acid residues | 70 | 74 | 73 | 98 | 80 |
| Ligands | Mg<sup>2+</sup>: 1 | Mg<sup>2+</sup>: 1 | Mg<sup>2+</sup>: 1 | Mg<sup>2+</sup>: 1 | Mg<sup>2+</sup>: 1 |
|  | Zn<sup>2+</sup>: 2 | Zn<sup>2+</sup>: 2 | Zn<sup>2+</sup>: 2 | Zn<sup>2+</sup>: 2 | Zn<sup>2+</sup>: 2 |
|  | CHAPSO: 4 | CHAPSO: 4 | CHAPSO: 3 | CHAPSO: 3 | CHAPSO: 4 |
| *B* factors (Å<sup>2</sup>) |  |  |  |  |  |
| Protein | 96.51 | 77.22 | 58.54 | 60.27 | 129.11 |
| Nucleic acids | 188.71 | 185.22 | 158.09 | 145.46 | 231.17 |
| Ligand | 84.01 | 70.81 | 71.23 | 64.42 | 117.25 |
| R.m.s. deviations |  |  |  |  |  |
| Bond lengths (Å) | 0.004 | 0.003 | 0.004 | 0.006 | 0.004 |
| Bond angles (°) | 0.591 | 0.516 | 0.562 | 0.628 | 0.643 |
| Validation |  |  |  |  |  |
| MolProbity score | 2.18 | 1.79 | 2.07 | 2.22 | 2.45 |
| Clashscore | 6.10 | 4.73 | 5.48 | 5.85 | 8.61 |
| Poor rotamers (%) | 3.93 | 1.79 | 3.39 | 4.33 | 4.57 |
| Ramachandran plot |  |  |  |  |  |
| Favored (%) | 94.28 | 94.92 | 94.54 | 93.68 | 92.0 |
| Allowed (%) | 5.62 | 5.08 | 5.4 | 6.26 | 7.79 |
| Disallowed (%) | 0.11 | 0.0 | 0.05 | 0.05 | 0.21 |

<sup>a</sup> All micrographs collected using a Titan Krios (ThermoFisher Scientific) microscope with an accelerating voltage of 300 kV.
<sup>b</sup> Nominal consensus map resolutions are from single particle reconstructions prior to Bayesian polishing in RELION <sup>73</sup>.

Examination of the four consensus maps revealed features indicative of structural heterogeneity in the nonconserved insert of σ^70^ (σ^70^_NCR_; σ^70^ residues 128-376), the sequence insertion in the trigger-loop (SI3; β’ 942-1131), the DNA upstream of the -35 element, and the σ-C-terminal domains (σCTDs). To isolate conformational changes involving the promoter -10 element and the RNAP active site cleft, we performed 3DVA within a mask encompassing the entrance to the cleft (σ^70^_2_ and σ^70^_3_ but excluding the σ^70^_NCR_), the pincers (βprotrusion, βlobe, and clamp), and the downstream end of the channel (including the β’jaw) (Extended Data Fig. 4).

Clamp opening/closing, a well-studied functional characteristic of cellular RNAPs ^28–35^ and a target of antibiotics ^36–41^, was a major mode of motion observed in all the 3DVAs (Fig. 1b). Analysis of these 3DVA trajectories revealed a correlation between the clamp motions and progression along the RPo formation pathway (Fig. 1c). We therefore focused on this component in the 3DVA analyses (Supplementary Video 2).

Each dataset (1_120ms_, 2_120ms_, 3_120ms_, and 4_500ms_) was analyzed independently, as were the combined particles from the 120 ms experiments (123_120ms_). Using masked 3DVA cluster analysis of the clamp open/close mode of the 120 ms datasets, we repeatedly found the same three distinct intermediates and sometimes found a fourth, poorly populated intermediate. With the 500 ms dataset, the same four intermediates were observed. We designated the intermediates I1a, I1b, I1c, and I1d to indicate that they are part of the ‘I1 ensemble’ that precedes the conversion to RPo (Fig. 1a) ^5^.

Due to the internal consistency of the independent analyses, we combined all the particles into one large dataset (123_120ms_4_500ms_) to increase the signal of any low-abundance states. The combined dataset was analyzed for structural heterogeneity using the same masked 3DVA procedure (Extended Data Fig. 4). This yielded the same four intermediates (I1a, I1b, I1c, I1d) observed in the analyses of the individual datasets (1_120ms_, 2_120ms_, 3_120ms_, and 4_500ms_). The fractions of particles distributed into each intermediate from each of the three independent 120 ms datasets were nearly the same (standard deviations of average particle fractions < 10%; Supplementary Table 1a), indicating that the tr-Spotiton device and the analysis pipelines were reproducible. The population distribution of dataset 4_500ms_ was also similar but significantly different according to Jensen-Shannon distances (Supplementary Table 1b)^42^, being skewed towards the most advanced intermediate I1d (Supplementary Table 1, Fig. 1c; Extended Data Fig. 5).

The cryo-EM maps derived from the combined dataset were used for model building and refinement (Table 1, Extended Data Figs. 6 and 7). The intermediate structures were ordered along the RPo formation pathway such that the DNA-Eσ^70^ interface area and the downstream boundary of the DNA-Eσ^70^ contacts increased, while the root-mean-square deviation of σ-carbon positions of each complex compared to σP_R_-RPo (7MKD)^7^ decreased (Fig. 1c). These metrics of progress along the RPo formation pathway correlated with the RNAP clamp position, with I1a having a relatively open clamp [6.9° compared to RPo (0°)] and the clamp closing between 1° to 2° in each subsequent intermediate (Figs. 1b and 1c). A control dataset to examine whether CHAPSO influenced these results was collected using *1H*, *1H*, *2H*, *2H*-perfluorooctyl)phosphocholine (fluorinated Fos-Choline-8, or FC8F, an alternative detergent discovered to mitigate particle orientation bias) and a mixing time of 500 ms. The same structural intermediates were observed (Extended Data Fig. 8).

### Duplex DNA rapidly unwinds atλP_R_

By 120 ms at RT, all λP_R_-Eσ^70^ complexes converted to intermediates in which DNA melting was nucleated (Fig. 2): the W-dyad was edge-on (forming a W-wedge), the -12 bp was open, and A_-11_(nt) was extra-helical (Fig. 3). The single-stranded t-strand within the transcription bubble was dynamic (no interpretable cryo-EM density) in all intermediates. We did not observe upstream DNA wrapping on Eσ^70^ in any I1 intermediate (DNA to -85; Extended Data Fig. 1a).

**Fig. 2.**
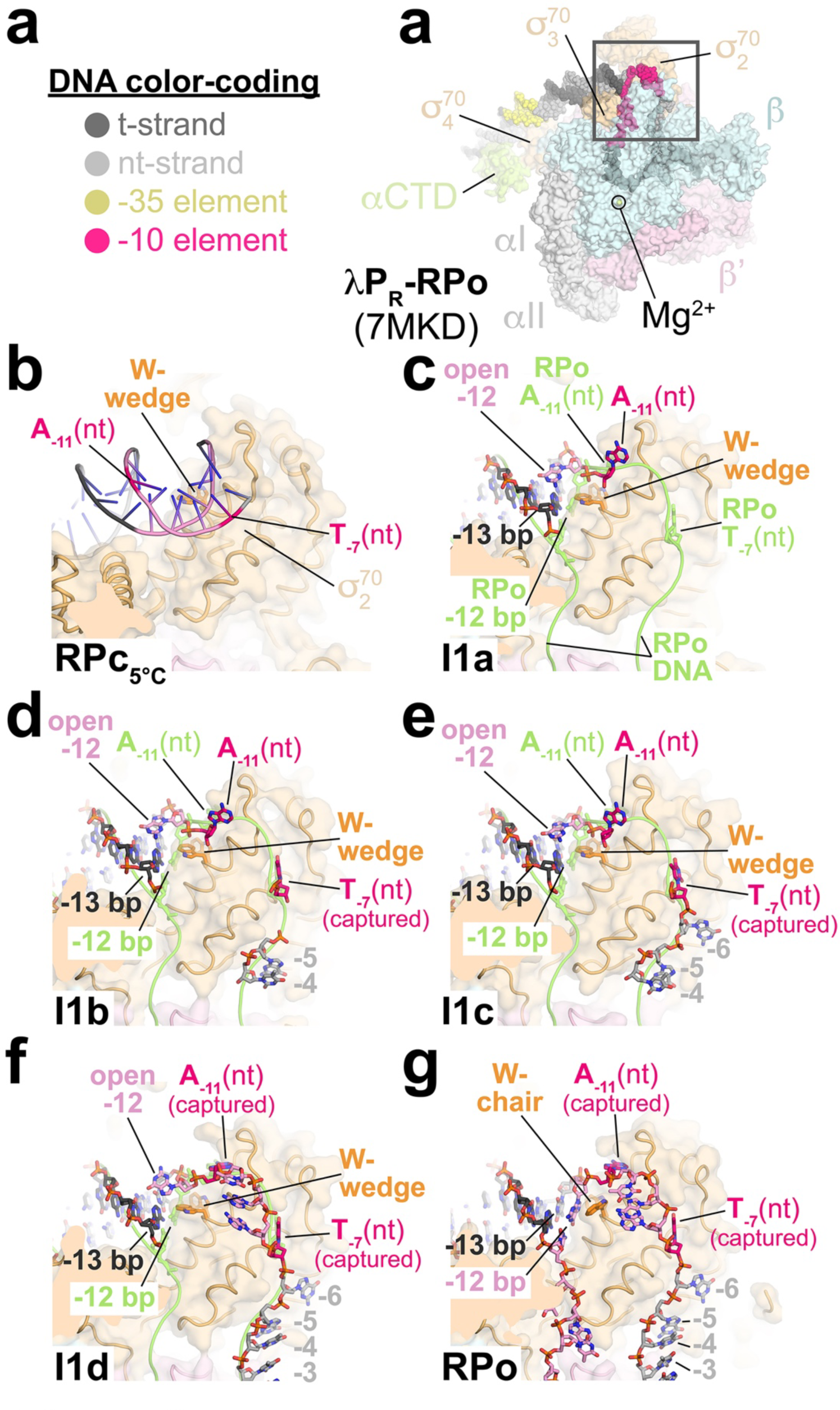
Transcription bubble nucleation. **a.** Top view of σP_R_-RPo (7MKD) ^7^. Eσ^70^ is shown as a transparent molecular surface. The DNA is shown as atomic spheres, color-coded as shown on the left. **b.** The boxed region in (**a**) is magnified (the β subunit is removed for clarity), showing the region of transcription bubble nucleation of RPc_5°C_. Protein is shown as a backbone worm with a transparent molecular surface. The side chains of the σ^70^ W-dyad (*Eco* σ^70^ W433/W434) are shown (W-wedge conformation). The duplex (closed) DNA is shown in cartoon format. **c.** As in (**b**) but showing I1a. The DNA is shown in stick format. The -12 bp is open due to steric clash with the W-wedge. A_-11_(nt) is flipped but not captured. For reference, the path of the DNA in RPo is shown in chartreuse, with the positions of key nucleotides shown in stick format [-12 bp, A_-11_(nt), T_-7_(nt)]. **d.** Magnified view of I1b. T_-7_(nt) is captured in its cognate σ^70^ pocket. **e.** Magnified view of I1c. **f.** Magnified view of I1d. A_-11_(nt) is captured but the W-dyad remains in the W-wedge conformation and the -12 bp remains open. **g.** Magnified view of RPo. The σ^70^ W-dyad has isomerized to the chair conformation, allowing repairing of the -12 bp. Also see Extended Data Figs. 2-11 and Supplementary Video 2.

**Fig. 3.**
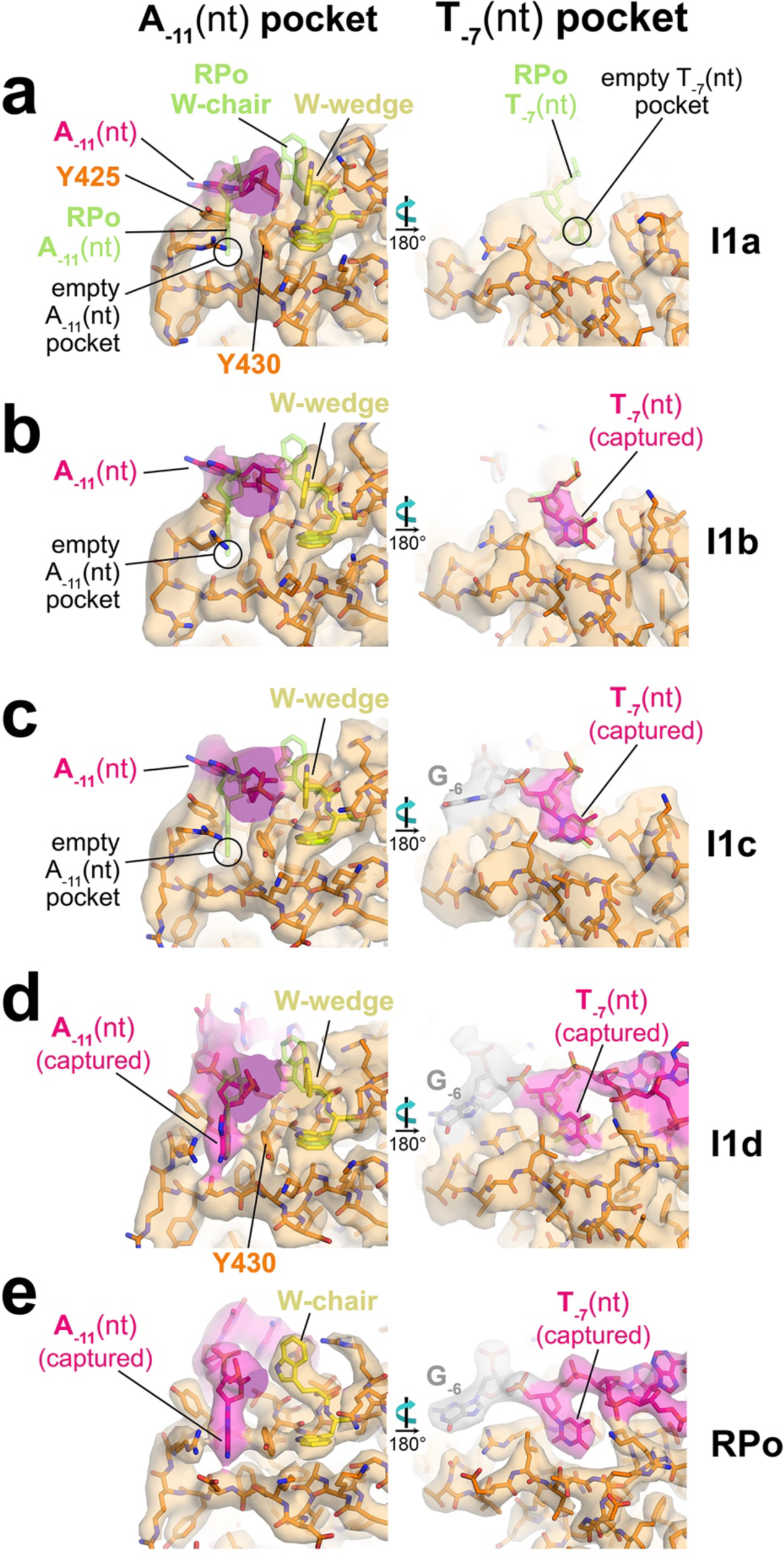
Capture of T_-7_(nt) before A_-11_(nt). Each panel shows two views of the σP_R_-Eσ^70^ complexes. σ^70^ and DNA (color-coded as in Fig. 3) are shown in stick format; carbon atoms are colored orange but the W-dyad is highlighted in yellow. Transparent cryo-EM density (local-resolution filtered ^75^) is superimposed. For reference, the positions of key RPo elements are shown in stick format and colored chartreuse [A_-11_(nt), T_-7_(nt), W-dyad in chair conformation]. (*left*) The σ^70^_2_-A_-11_(nt) pocket, viewed from upstream. (*right*) The σ^70^-T_-7_(nt) pocket. **a.** I1a; (*left*) A_-11_(nt) is flipped but not captured [the σ^70^_2_-A_-11_(nt) pocket is empty] and the W-dyad is in the wedge (edge-on) conformation, (*right*) The T_-7_(nt) pocket is empty. **b.** I1b; (*left*) A_-11_(nt) is flipped but not captured [the σ^70^_2_-A_-11_(nt) pocket is empty] and the W-dyad is in the wedge (edge-on) conformation, (*right*) T_-7_(nt) is captured. **c.** I1c; (*left*) A_-11_(nt) is flipped but not captured [the σ^70^_2_-A_-11_(nt) pocket is empty] and the W-dyad is in the wedge conformation, (*right*) T_-7_(nt) is captured. **d.** I1d; (*left*) A_-11_(nt) is captured but the W-dyad is still in the wedge conformation, (*right*) T_-7_(nt) is captured. **e.** RPo; (*left*) A_-11_(nt) is completely captured and the W-dyad is in the chair conformation, (*right*) T_-7_(nt) is captured. Also see Extended Data Fig. 11.

In I1a, I1b, and I1c, cryo-EM density features for σ^70^_1.1_ were present in the RNAP cleft (Fig. 4). In I1b and I1c, the single-stranded nt-strand gradually appears (I1b, -7, -5, -4; I1c, -7, -6, -5, -4; Figs. 2d and 2e), following approximately the same path as in RPo (Figs. 2a and 2g). At I1d, additional large scale changes ocurred: the DNA strands were unambiguously unwound from -12 to +2, A_+1_(nt) stacked on βW183 (as in RPo), and double-stranded DNA (from +3 to +10 visualized) occupied the RNAP downstream channel in place of σ^70^_1.1_ (Fig. 4e).

**Fig. 4.**
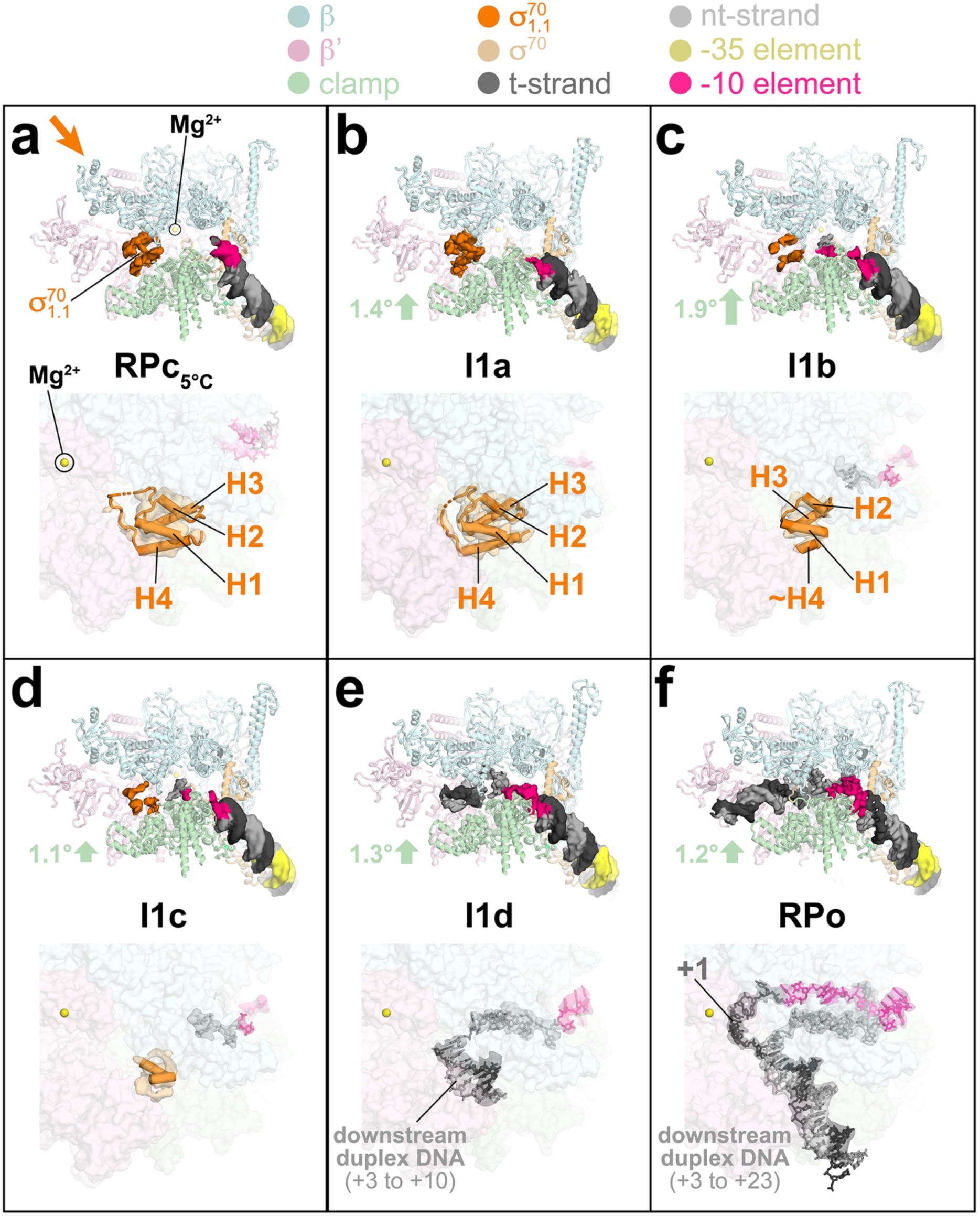
RNAP clamp closure partially unfolds and ejects σ^70^_1.1_. Each panel shows two views of the σP_R_-Eσ^70^ complexes (σ^70^_NCR_ omitted for clarity), with color-coding shown at the top. (*top*) View into the RNAP active-site cleft. Eσ^70^ is shown as a backbone cartoon. Cryo-EM density for σ^70^_1.1_ (orange) and the DNA are also shown. (*bottom*) View focusing on σ^70^_1.1_ [viewed from the direction of the thick orange arrow in (**a**)]. Eσ^70^ is shown as a transparent molecular surface, but σ^70^_1.1_ is shown as a backbone cartoon with cylindrical helices. DNA is shown in stick format. Transparent cryo-EM density (local-resolution filtered ^75^) for σ^70^_1.1_ (orange) and the DNA is also shown. **a.** RPc_5°C_. **b.** I1a. Closure of the clamp from the previous intermediate is denoted by the thick green arrow. **c.** I1b; σ^70^_1.1_-H4 becomes largely disordered. **d.** I1c; cryo-EM density for σ^70^_1.1_ is present but becomes mostly uninterpretable. **e.** I1d; σ^70^_1.1_ is replaced by duplex DNA (+3 to +10) in the RNAP cleft. **f.** RPo. Also see Extended Data Figure 12.

### Intermediates populated at equilibrium at low temperature at σP_R_

The observation of full transcription bubble formation in the I1 ensemble (I1d; Fig. 4e) was unexpected ^5,25,43,44^. To confirm the structural characteristics of the I1 ensemble as observed by tr-Spotiton at RT, we examined σP_R_-Eσ^70^ complexes trapped at equilibrium at 5°C using standard cryo-EM grid preparation methods. On σP_R_ at temperatures below 7°C, the high activation energy (E_a_ ∼34 kcal) to reach the transition state between I1 and I2 blocks conversion to complexes past I1, leaving only I1 and any preceding complexes at equilibrium ^25,44^.

σP_R_-Eσ^70^ complexes were pre-formed on ice and then spotted and blotted on grids held at 5°C. After collection and processing of 2764 micrographs using the same pipeline as the tr-Spotiton datasets (Table 1; Extended Data Fig. 9), we identified an earlier intermediate not seen in the RT tr-spotiton experiments (RPc_5°C_, nominal resolution 3.1 Å, Table 1; Extended Data Figs. 9 and 10a-d), and two complexes that share similarities with I1c and I1d, with population distribution RPc_5°C_, 52%; I1c_5°C_, 21%; I1d_5°C_, 27% (Extended Data Figs. 9 and 10e-m). These data are consistent with our finding that the I1 ensemble contains a significant fraction of DNA-Eσ^70^ complexes with a fully-melted transcription bubble.

### Transcription bubble nucleation: -12 bp opening and capture of T_-7_(nt) before A_-11_(nt)

In the earliest intermediates detected at σP_R_ at RT (I1a, I1b, I1c), the transcription bubble was nucleated; A_-11_(nt) was flipped but not yet captured in the σ^70^ pocket (Figs. 2c-e, 3a-c). Capture of A_-11_(nt) did not occur until I1d (Figs. 2f and 3d). As early as I1b we observed clear cryo-EM density for T_-7_(nt) capture (Fig. 3b). Thus, T_-7_(nt) capture preceded A_-11_(nt) capture ^45^, illuminating why T_-7_(nt) is overrepresented in fast melting -10 element sequences ^46^.

Strikingly, through all the intermediates observed here, from initial transcription bubble nucleation (I1a) to full bubble formation (in I1d), the σ^70^ W-dyad remained in its edge-on (wedge) conformation (Figs. 2b-f and 3a-d). As a result, in all the nucleated intermediates (I1a-I1d), the -12 bp was disrupted by steric clash with the edge-on W-wedge (Figs. 2c-f; Extended Data Fig. 11). The un-paired -12 and -11 nt-strand bases were essentially exposed to solution and highly dynamic, although the A_-11_(nt) base may stack on σ^70^-Y425 in I1a-c (Fig. 3a-c).

### Stepwise closure of the RNAP clamp partially unfolds and ejects σ^70^

As the clamp closes through the RPo formation pathway, a striking feature emerges: σ^70^ becomes progressively disordered. In RPc_5°C_ and I1a, the four σ-helices of σ^70^ occupy the RNAP cleft (H1-H4, Figs. 4a and 4b) as seen in Eσ^70^ ^16^. In the conversion from I1a to I1b, cryo-EM density for much of H4 and for the 37-residue “linker” that connects H4 _1.2_ (*Eco* σ residues 75-91) disappears (Fig. 4c). Upon further closure of the clamp _70 70_ in I1c, the cryo-EM density for σ^70^ further fragments and becomes largely uninterpretable except for two tube-like densities modeled as σ-helices (Fig. 4d). The σ^70^ finally disappears in I1d (Figs. 4e).

To test the importance of σ^70^ unfolding for RPo formation, we introduced double-cysteine substitutions into σ^70^ (in the background of a Cys-less σ^70^ derivative) ^47^ designed to form interhelical disulfide bonds under oxidizing conditions to interfere with σ^70^ unfolding (Fig. 5a). Three Cys-pairs were constructed: Q8C-P32C

**Fig. 5.**
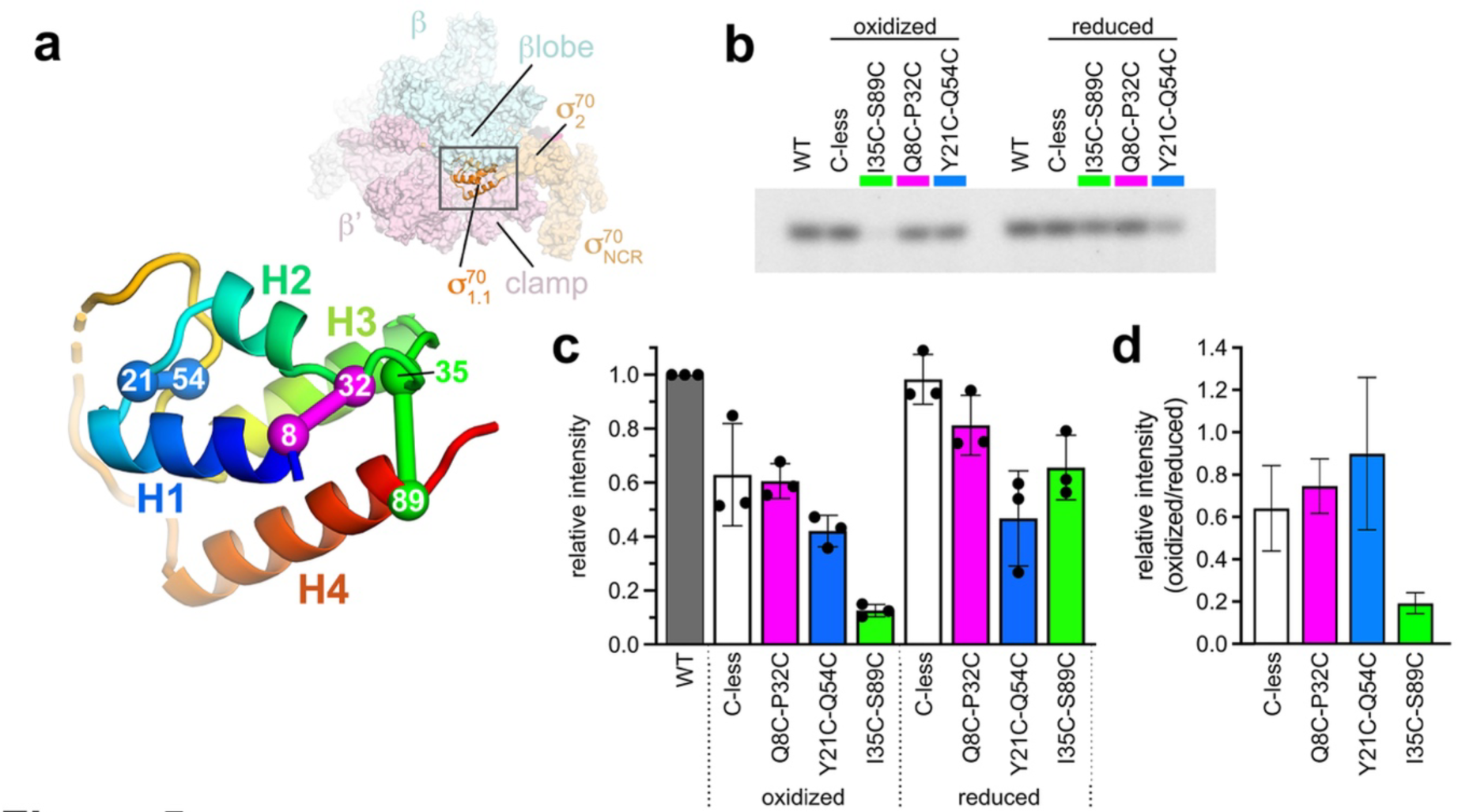
σ^70^_1.1_-H4 unfolding is necessary for RPo formation. **a.** (*top*) Overall view of I1a. Eσ^70^ is shown as a transparent molecular surface but σ^70^_1.1_ (orange) is shown as a backbone cartoon. (*bottom*) σ^70^_1.1_ from boxed region above, colored as a rainbow ramp from the N-terminus (blue) to the C-terminus of H4 (red). Pairs of residues substituted with Cysteine (in the background of a Cysteine-less σ^70^ ^47^) are shown as Cσ spheres, with engineered disulfide bonds illustrated with thick lines. **b.** Synthesis of abortive products (ApUp*G, where * denotes σ-[P^32^]) from σP_R_ by Eσ^70^’s containing WT or mutant σ^70^’s as indicated. Oxidizing conditions (to form disulfide bonds, left) are compared with reducing conditions (right). **c.** Relative intensity of abortive products (normalized with respect to WT σ^70^) for mutant σ^70^’s under oxidizing and reducing conditions. Error bars denote the standard deviation of n=3 measurements. **d.** Ratio of abortive product intensity for oxidized/reduced conditions [error bars were calculated by error propagation from the standard deviations shown in (**c**)]. The I35C-S89C disulfide, crosslinking σ^70^_1.1_-H4 to H3, is severely defective under oxidizing conditions. Also see Extended Data Fig. 12.

(crosslinking H1 to H3), Y21C-Q54C (crosslinking the H1-H2 linker with the H3-H4 linker), and I35C-S89C (crosslinking the H3 with H4). Each derivative had a higher mobility under oxidizing conditions than reducing conditions when analyzed by denaturing polyacrylamide gel electrophoresis (Extended Data Fig. 12a), indicating that the expected disulfide bonds formed. Comparison of abortive initiation transcription activity under oxidizing vs. reduced conditions revealed that the I35C-S89C (H3-H4) crosslink was severely defective in producing a transcript (Figs. 5b-d), suggesting that RPo cannot form. This is consistent with the order-to-disorder transition of σ^70^_1.1_-H4, begining at the I1a -> I1b transition, being essential for progress through the RPo formation pathway.

## Discussion

We used cryo-EM to determine high-resolution structures of early RPo formation intermediates at σP_R_ (Fig. 6). Early intermediates were trapped at equilibrium at low temperature (RPc_5°C_), or at RT in real time (non-equilibrium) using tr-Spotiton (Extended Data Fig. 1b) ^20^. The structures delineate conformational changes in both Eσ^70^ and promoter DNA on the pathway to forming RPo and reveal unanticipated features. Analysis of the structures of early RPo intermediates allows direct comparison with extensive mechanistic studies of DNA opening on σP_R_ ^5^ and provides unprecedented insights into the mechanism of transcription bubble nucleation (Fig. 2) and σ^70^ ejection from the RNAP cleft (Figs. 4 and 5). This study identifies early intermediates of RPo formation at one promoter, σP_R_, but the invariant Eσ^70^ architecture and conserved nature of promoter -10 and -35 elements ^4^ suggests that these structures define key steps of DNA opening at most Eσ^70^ promoters ^10^.

**Fig. 6.**
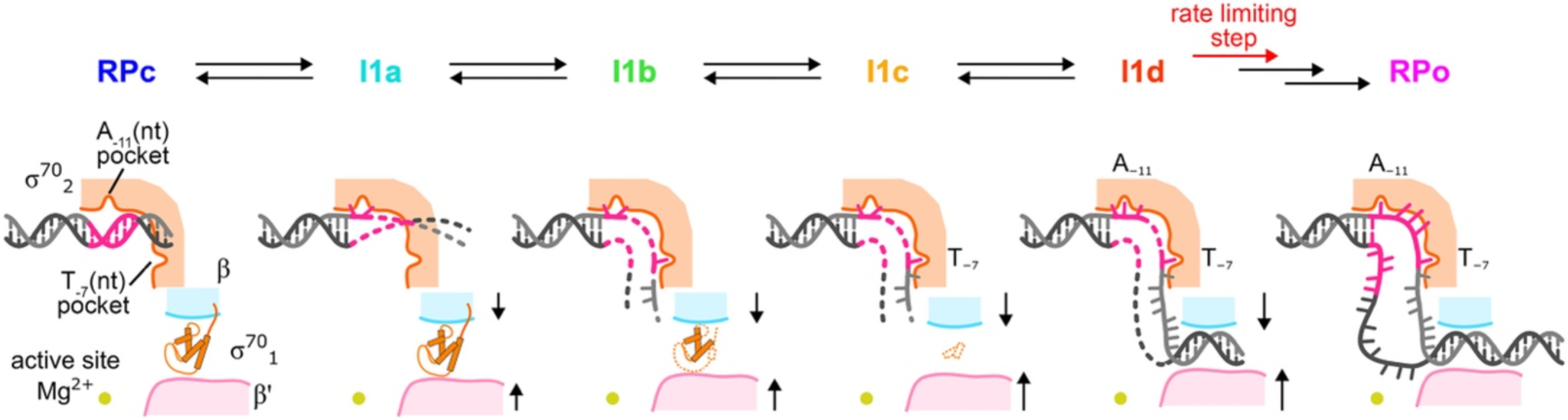
Schematic overview of initial steps of promoter opening at σP_R_. The region of transcription bubble nucleation is shown for RPc, each I1 intermediate, and RPo. The RNAP active-site Mg^2+^ is shown as a yellow sphere. The σ^70^ domain, with its A_-11_(nt) and T_-7_(nt) pockets, and σ^70^, are shown in orange. In RPc to I1c, elements of σ^70^ are in the RNAP active-site cleft between elements of β (cyan) and β’ (pink). Closure of the clamp is denoted by the black arrows. The DNA is shown as a backbone worm (-10 element colored hot pink). Poorly-resolved regions of the DNA or _70 1.1_ are illustrated by dashed lines.

### A closed complex observed at 5°C (RPc_5°C_)

An RPc was first observed by cryo-EM with *Eco* Eσ^70^ at the *rpsT*P2 promoter (6PSQ) where stabilization of early intermediates by the transcription factor TraR likely populated RPc ^10^. RPc on σP_R_ was populated in samples prepared at equilibrium at low temperature (RPc_5°C_; Figs. 2b and 4a) but not in any of the tr-Spotiton datasets (prepared at RT). At the relatively high final concentrations of Eσ^70^ and DNA on the tr-Spotiton grids (15-30 μM), we expect RPc to fully occupy the promoter DNA and convert to I1 in <10 ms ^5,27^.

The *rpsT*P2-RPc and σP_R_-RPc_5°C_ have very similar structural features; the promoter DNA is completely duplex and outside the RNAP cleft, the RNAP clamp is relatively open (*rpsT*P2-RPc, 8.5°; σP_R_-RPc_5°C_, 6.9°, using σP_R_-RPo as 0° reference), and there are no Eσ^70^-DNA contacts downstream of -2 nor base-specific contacts in the promoter -10 element ^10^. A superposition of the RNAP structures (excluding the βlobe-Si1, which is repositioned due to TraR binding in *rpsT*P2-RPc) yielded a root-mean-square-deviation (rmsd) of 1.2 Å over 2,084 RNAP σ-carbons. An RPc with *Mycobacterium tuberculosis* RNAP with the transcription factor WhiB7 on the p*WhiB7* promoter shares these same structural characteristics ^48^. Based on these observations, the most straightfoward proposal is that this RPc structure is the first, or one of the first, promoter DNA-RNAP complexes at most, if not all, Eσ^70^ promoters.

### The RNAP clamp

Clamp dynamics play an important role in promoter melting for all cellular RNAPs ^10,19,32,49–51^. Several antibiotics, including clinically important Fidaxomicin ^39–41^, bind RNAP in the switch regions (essentially hinges for the clamp motion) and interfere with initiation, highlighting the importance of clamp dynamics ^38^. In 3DVAs of the heterogeneous σP_R_-Eσ^70^ conformations observed in the tr-Spotiton samples, clamp opening and closing was the major mode of motion that correlated with other structural features expected to be associated with progress on the RPo formation pathway (Supplemental Video 2). As the clamp closed from RPc_5°C_ (6.9° open) to RPo (0°), the DNA-Eσ^70^ contacts progressed downstream while the DNA-Eσ^70^ interface area increased, and the RNAP conformation became more RPo-like (Fig. 1c).

We propose that entry of the DNA into the RNAP cleft and gradual establishment of in-cleft DNA-RNAP interactions (Fig. 2) drive gradual clamp closure (Fig. 1b). The clamp closure in turn drives many of the conformational changes that propel the complex along the RPo formation pathway. As DNA melting propagates downstream after nucleation at -12/-11, stepwise clamp closing appears to position Eσ^70^ structural elements to establish sequence-specific interactions with the -10 hexamer and discriminator ^52–56^ promoter elements. Perhaps most strikingly, for σP_R_, interactions with nt-strand discriminator bases (-6 to -4) appear to occur along with capture of T_-7_(nt), followed by A_-11_(nt) (Figs. 2d, 2e, and 6). Early establishment of discriminator interactions is consistent with the critical role of the discriminator in transcription start site selection ^56^. We note that DNA sequence differences in these promoter elements (which will affect the lifetimes and relative populations of melting intermediates) may alter the order in which they are stabilized by interactions with Eσ^70^. Promoter sequence effects on the distribution of melting intermediates provides opportunities for promoter-specific regulation by factors that don’t bind DNA, such as DksA ^57^

### Clamp closure drives coupled unfolding and extrusion of σ^70^ out of the RNAP cleft

Clamp closure appears to drive the coupled unfolding and extrusion of σ^70^ out of the RNAP cleft (Fig. 4) by a mechanism similar to protein unfolding induced by high hydrostatic pressure ^58^. As the clamp closes, the volume available for σ^70^, located between the closing RNAP pincers, decreases. In response, elements of σ^70^ unfold and extrude out the cleft, decreasing the volume occupied by σ^70^ inside the cleft. σ^70^ -H4, known to be semi-stable and located at the mouth of the cleft ^16^, and H3-H4 linker were squeezed out of the cleft by the closing clamp. The unfolding and extrusion _70 1.1_-H4 appears to be essential for further progress on the RPo formation pathway (Fig. 5); crosslinking of H4 to H3, preventing unfolding of H4, may inhibit further closure of the clamp which would be required to completely dislodge σ^70^ and establish the necessary promoter DNA-RNAP contacts.

### Clamp closure drives transcription bubble nucleation

The σ^70^-W-dyad remained edge-on in all the intermediates observed here (RPc_5°C_, I1a, I1b, I1c, I1d), in common with the earliest intermediates at *rpsT*P2 (RPc and T-RPi1) ^10^. Therefore, we propose that this W-dyad conformer may be a shared feature of early intermediates at most Eσ^70^ promoters. The edge-on W-dyad comprises a stable structural unit ^59^ that may act as a wedge (the W-wedge) to nucleate the transcription bubble. Clamp closure may squeeze the DNA between the βprotrusion and σ^70^_2_, driving the DNA into the σ^70^-W-wedge and sterically disrupting the -12 and -11 bps, nucleating the transcription bubble (Fig. 2).

Opening of the -12 bp due to steric clash with the W-wedge was first observed in an early intermediate at *rpsT*P2, T-RPi1 ^10^. This was a surprising observation since the - 12 position [immediately upstream of the critical A_-11_(nt) base] is base-paired in all known RPo structures, including *rpsT*P2-RPo andλP_R_-RPo ^7,10^. Thus, at least at these two promoters, nucleation disrupts the -12 bp early in RPo formation. Our finding of - 12 bp opening in all I1 ensemble intermediates (Fig. 2) suggests that unpairing and re-pairing at -12 may be a conserved feature of RPo formation at all σ^70^-family promoters and could explain the conservation of T_-12_(nt) within the -10 element ^4^.

The W-dyad remained in the wedge conformation even in I1d, where full promoter opening was clearly achieved (Fig. 2f). This observation contrasts with intermediates at *rpsT* P2, where the W-wedge was observed only in the earliest nucleated intermediate, T-RPi1; the W-dyad isomerized to the chair conformation in subsequent intermediates even though full promoter melting was not achieved ^10^. The *rpsT*P2 intermediates were trapped and prepared for cryo-EM at equilibrium (i.e. after long incubation times) while theλP_R_ I1 intermediates observed here were prepared for cryo-EM in real-time at non-equilibrium conditions by tr-Spotiton after 120 or 500 ms mixing times. We suggest that during RPo formation, W-dyad isomerization from the wedge to the chair conformation occurs at a slow rate compared to full bubble opening; W-dyad isomerization may have occurred in the *rpsT* P2 intermediates (trapped by TraR and, in some cases, promoter mutations) due to the long incubation times and equilibrium conditions ^10^. This highlights a major advantage of time-resolved approaches for cryo-EM sample preparation, critical for trapping important conformations of macromolecular complexes in transient, non-equilibrium states.

### Full promoter opening precedes the rate-limiting step atλP_R_

Before recent cryo-EM studies ^10,19^, descriptions of RPo intermediates were largely obtained by chemical or enzymatic footprinting of complexes trapped at equilibrium under perturbing solution conditions (low temperature, inhibitors) ^60–63^. Permanganate ion (MnO_4_^-^) oxidation of unstacked thymines was used to detect DNA melting ^64^. Protection of the DNA backbone from cleavage by DNase I or hydroxyl-radicals was used to map RNAP-DNA interactions ^65,66^.

Footprinting of Eσ^70^-λP_R_ complexes equilibrated at 0°C (so comprising the I1 ensemble) did not detect any unstacked nt-strand thymines despite DNA backbone protection extending to ∼+20 ^44^. To rationalize these results, the DNA in I1 was proposed to be sharply bent or kinked at -11/-12, directing the duplex DNA into the RNAP cleft ^44^. Subsequent studies found that the rate constants for the appearance of oxidized thymines were similar to those for the conversion of I1 to I2 ^27^. These and other studies led to the conclusion that I1 was closed (no bubble nucleation), and that the rate-limiting step for RPo formation atλP_R_ (the I1 -> I2 conversion) corresponded with the formation of the transcription bubble ^67^.

By the time I1d forms, thymines on the nt-strand are stacked (-4, -3) or bound in a pocket (-7). Lacking stabilizing RNAP interactions (and thus lacking a driving force for unstacking), t-strand bases may remain largely stacked in I1. Although the rates of interconversion between the I1 ensemble intermediates are unknown, they are likely relatively fast: in the absence of protein-DNA interactions, lifetimes of transiently open DNA bubbles (2-10 bp) ^68^ or extrahelical bases ^69^ range from 10^-2^ to 10^-6^ s. Because the MnO_4_^-^ oxidation reaction rate constant is relatively slow (∼15 M^-1^ s^-1^) and decreases with decreasing temperature ^70^, we suggest that the footprinting conditions (low temperature, low MnO_4_^-^ dose) used for σP_R_ may have prevented detection of DNA opening in I1.

In support of this hypothesis, fast MnO_4_^-^ and hydroxyl-radical footprinting studies at 37°C on the T7 A1 promoter revealed that DNA melting preceded the rate-limiting step and the formation of stable downstream contacts ^71^. Mechanistic studies of RPo formation at a consensus promoter ^72^ and at gal P1 ^45^ found that the rates of unquenching (unstacking) 2-amino purine fluorescence were faster than the rates of forming RPo. These results are consistent with our findings from time-resolved cryo-EM on σP_R_ that full promoter opening precedes the rate-limiting step for RPo formation.

### Rate-limiting steps in RPo formation

If neither σ^70^ ejection nor DNA opening are rate-limiting at σP_R_, what is? Conversion from I1 to I2 (Fig. 1a) requires overcoming a large positive activation energy ^25^. Once I2 forms, thymines on the t-strand become solvent-accessible; T_+1_(t) is as MnO_4_^-^-reactive as in RPo ^67^. These results, along with the observation that the σ^70^-finger in the I1 ensemble clashes with the position of the t-strand in RPo (Extended Data Fig. 12b) suggest that the rate-limiting step at σP_R_ loads the t-strand in the active site by repositioning the σ^70^-finger and unstacking t-strand bases. Other potentially rate-limiting changes occur as I1 converts to I2 and then RPo: an intricate network of interactions form between σ^70^ the β “gate loop”, and the nt-strand discriminator bases ^7^, and the W-wedge isomerizes to “chair”, allowing the bp at - 12 to re-pair (Fig. 3d). Further kinetic-mechanistic and cryo-EM studies will be required to tease apart these conformational changes and how they are driven by promoter DNA sequence.

## Acknowledgments

We thank E.A. Campbell, R. Gourse, D. Jensen, R. Landick, W. Ross, and members of the Darst-Campbell laboratory for helpful discussions. Some of the work reported here was conducted at the Simons Electron Microscopy Center (SEMC) and the National Resource for Automated Molecular Microscopy (NRAMM) located at the New York Structural Biology Center, supported by grants from the NIH National Institute of General Medical Sciences (P41 GM103310), NYSTAR, and the Simons Foundation (SF349247). A.U.M. received support through a fellowship from the Swiss National Science Foundation. A.U.M. is an Agouron Institute Awardee of the Life Sciences Research Foundation. This work was supported by NIH grant R35 GM118130 and Re-Entry Supplement R35 GM118130-04S1 to S.A.D.

## Author contributions

Conceptualization; R.M.S, A.U.M., J.C., B.M., C.S.P., B.C., S.A.D.; Protein purification, biochemistry; R.M.S, A.U.M., J.C., B.M.; Cryo-EM specimen preparation; R.M.S., A.U.M., J.C., B.M., W.C.B., V.P.D.; Cryo-EM data collection and processing: R.M.S., A.U.M., J.C., B.M., V.P.D., K.M., J.H.M., E.T.E., L.Y.Y.; Model building and structural analysis: R.M.S., A.U.M., B.M., S.A.D.; Funding acquisition and supervision: C.S.P., B.C., S.A.D.; Manuscript first draft: R.M.S., A.U.M., S.A.D.; All authors contributed to finalizing the written manuscript.

## Methods

No statistical methods were used to predetermine sample size. The experiments were not randomized, and the investigators were not blinded to allocation during experiments and outcome assessment.

### Protein expression and purification

Eco *core RNAP* [σ_2_ββ’(His)_10_ω] was purified largely as described previously ^33^. A pET-based plasmid overexpressing each subunit of RNAP (full-length σ, β, ω) as well as β’-PPX-His_10_ (PPX; PreScission protease site, LEVLFQGP, Cytiva, Marlborough, MA) were co-transformed with a pACYCDuet-1 plasmid expressing *Eco rpo*Z into *Eco* BL21(DE3). The cells were grown in the presence of 100 µg/mL ampicillin, 34 μg/mL chloramphenicol, and 0.5 mM ZnCl_2_ to an OD_600_ of 0.6 in a 37°C shaker. Protein expression was induced with 1 mM IPTG (final concentration) for 4 hours at 30°C. Cells were harvested by centrifugation and resuspended in lysis buffer [50 mM Tris-HCl, pH 8.0, 5% glycerol (v/v), 10 mM DTT, 1 mM PMSF, and 1x protease inhibitor cocktail (Sigma Aldrich)]. After French Press lysis at 4°C, the lysate was centrifuged twice (33,000 x g) for 30 minutes each. Polyethyleneimine [PEI, 10% (w/v), pH 8.0, Acros Organics - ThermoFisher Scientific, Waltham, MA] was slowly added to the supernatant to a final concentration of ∼0.6% (w/v) PEI with continuous stirring. The mixture was stirred at 4°C for an additional 25 min, then centrifuged for 1.5 hours (33,000 x g) at 4°C. The pellets were washed three times in lysis buffer + 500 mM NaCl. For each wash, the pellets were resuspended using a Dounce homogenizer, then centrifuged again. RNAP was eluted by washing the pellets three times with lysis buffer + 1 M NaCl. The PEI elutions were combined and precipitated with ammonium sulfate overnight. The mixture was centrifuged and the pellets were resuspended in RNAP buffer [20 mM Tris-HCl, pH 8.0, 5% glycerol (v/v), 5 mM DTT] + 1 M NaCl. The mixture was loaded onto three 5 mL HiTrap IMAC HP columns (Cytiva) for a total column volume (CV) of 15 ml. RNAP(β’-PPX-His_10_) was eluted with RNAP buffer + 250 mM imidazole. The eluted RNAP fractions were combined and dialyzed against RNAP buffer + 100 mM NaCl. The sample was then loaded onto a 35 mL Biorex-70 column (Bio-Rad, Hercules, CA), washed with 10 mM Tris-HCl, pH 8.0, 0.1 mM EDTA, 5% glycerol (v/v), 5 mM DTT, in a gradient from 0.2 M to 0.7 M NaCl. The eluted fractions were combined, concentrated by centrifugal filtration, then loaded onto a 320 mL HiLoad 26/600 Superdex 200 column (Cytiva) equilibrated in gel filtration buffer [10 mM Tris-HCl, pH 8.0, 0.1 mM EDTA, 0.5 M NaCl, 5% glycerol (v/v), 5 mM DTT]. The eluted RNAP was supplemented with glycerol to 23% (v/v), flash frozen in liquid N_2_, and stored at -80°C.

Eco *His_10_-SUMO-σ*^70^ was expressed and purified as described previously ^7^. Plasmids encoding *Eco* His_10_-SUMO-σ^70^ were transformed into *Eco* BL21(DE3) by heat shock. The cells were grown in the presence of 50 µg/mL kanamycin to an OD_600_ of 0.4 at 37°C, then the temperature was lowered to 30°C. At OD 0.6, protein expression was induced with 1 mM IPTG (final) for 2 hours. Cells were harvested by centrifugation and resuspended in sigma lysis buffer [20 mM Tris-HCl, pH 8.0, 5% glycerol (v/v), 500 mM NaCl, 0.1 mM EDTA, 5 mM imidazole] and flash frozen in liquid N_2_. Cells were then thawed on ice, 2-mercaptoethanol (BME) and PMSF were added to 0.5 mM and 1 mM, respectively. After French Press lysis at 4°C, cell debris was removed by centrifugation. The lysate was loaded onto two 5 mL HiTrap IMAC HP columns (Cytiva) for a total CV of 10 ml. His_10_-SUMO-σ^70^ was eluted at 250 mM imidazole in 20 mM Tris-HCl, pH 8.0, 500 mM NaCl, 0.1 mM EDTA, 5% glycerol (v/v), 0.5 mM BME. Peak fractions were combined, cleaved with Ulp1, and dialyzed against 20 mM Tris-HCl, pH 8.0, 500 mM NaCl, 0.1 mM EDTA, 5% glycerol (v/v), 0.5 mM BME, resulting in a final imidazole concentration of 25 mM. The sample was loaded onto one 5 mL HiTrap IMAC HP column (Cytiva) to remove His_10_-SUMO-tag along with any remaining uncut His_10_-SUMO-σ^70^. Tagless σ^70^ was collected in the flowthrough and concentrated by centrifugal filtration. Pooled, cleaved samples were diluted to 200 mM NaCl using buffer A [10 mM Tris-HCl, pH 8.0, 0.1 mM EDTA, 5% glycerol (v/v), 1 mM DTT] and loaded onto three 5 mL HiTrap Heparin columns (Cytiva) equilibrated in buffer A. The σ^70^ was eluted over a NaCl gradient from 200 mM to 1 M. Peak fractions were pooled, concentrated and buffer exchanged into Superdex equilibration buffer [20 mM Tris-HCl, pH 8.0, 0.5 M NaCl, 5% glycerol (v/v), 1 mM DTT] using centrifugal filtration (Amicon Ultra, MilliporeSigma, Burlington, MA). Concentrated sample was then loaded onto a HiLoad 16/60 Superdex 200 size exclusion column (Cytiva); peak fractions of σ^70^ were pooled, supplemented with glycerol to a final concentration of 20% (v/v), flash-frozen in liquid N_2_, and stored at -80°C.

### Preparation of Eσ^70^ and σP_R_ DNA for cryo-EM

Eσ^70^ was formed by mixing core RNAP [σ_2_ββ’(His)_10_μ] with a 2-fold molar excess of σ^70^, and then incubating for 20-30 minutes in a heat block at 37°C. Eσ^70^ was separated from free σ^70^ on a Superose 6 Increase 10/300 GL size exclusion column (Cytiva) in gel filtration (GF) buffer (40 mM Tris-HCl, pH 8.0, 120 mM KCl, 10 mM MgCl_2_, 10 mM DTT). Peak fractions of the eluted Eσ^70^ were concentrated to ∼10-15 mg/mL by centrifugal filtration (Amicon Ultra, MilliporeSigma). Eσ^70^ samples were either taken on ice to the New York Structural Biology Center for same day tr-Spotiton experiments or flash frozen in N_2_(l) and stored at -80°C for later experiments.

All σP_R_ DNA constructs were commercially synthesized (Integrated DNA Technologies, Coralville, IA). To investigate upstream DNA-Eσ^70^ interactions formed at early times (tr-Spotiton experiments), σP_R_ oligomers from -85 to +20 (Extended Data Fig. 1a) ^76^ were used. For grids prepared using the Mark IV Vitrobot (equilibrium, 5.2 °C; Extended Data Fig. 9), σP_R_ DNA was -60 to +30 (Extended Data Fig. 9a) ^7^. For either oligomer, nt-strand and t-strand DNA (Extended Data Figs. 1a, 9a) was resuspended in annealing buffer (10 mM Tris-HCl, pH 8.0, 50 mM KCl, 0.1 mM EDTA). Equimolar amounts of the strands were mixed and incubated in a 95°C heat block for 5-10 minutes. The heat block was then removed to the benchtop where annealed strands slow cooled to room temperature. Annealed DNA was stored at -20°C before use.

### Cryo-EM grid preparation

*tr-Spotiton.* On-grid mixing experiments were performed using the tr-Spotiton instrument as described previously ^20,77^. Eσ^70^ and σP_R_ DNA were thawed on ice and diluted to 26 or 29.25 μM Eσ^70^ or to 52 or 58 μM σP_R_ DNA using GF buffer (1 Eσ^70^:2 DNA molar ratio). Before spraying, nanowire self-blotting grids ^22,78^ were plasma-treated (Gatan Solarus) at 5 W in H_2_ (g) and O_2_ (g) for 1 to 2 minutes. Grids were then placed in the tr-Spotiton chamber at RT and held at a relative humidity of 70 to 100%. Prior to loading into the dispensing tips, CHAPSO [(3-([3-cholamidopropyl]dimethylammonio)-2-hydroxy-1-propanesulfonate), Anatrace, Maumee, OH) 8 mM final, all 120 ms experiments and one 500 ms experiment] or fluorinated fos-choline-8 (FC8F; (1H, 1H, 2H, 2H-perfluorooctyl)-phosphocholine, Anatrace, Maumee, OH. 1.5 mM final, one 500 ms experiment) was added to each sample at RT (23 – 24 °C). After loading, equivalent streams of ∼50 pL droplets of σP_R_ DNA and Eσ^70^ were sequentially applied within 10 ms onto the grid (Extended Data Fig. 1b). Grid acceleration, deceleration, and final velocity parameters were chosen to achieve on-grid mixing times of 120 ms and 500 ms prior to the plunge into ethane (*l*). (See Supplemental Video 1).

*5°C equilibrium grids.* After mixing at RT, Eσ^70^ (15 μM final) and σP_R_ DNA (18 μM final) were equilibrated on ice in GF buffer and 8 mM CHAPSO. C-flat holey carbon grids (CF-1.2/1.3-4Au, Protochips, Morrisville, NC) were glow-discharged using a Solarus Plasma Cleaner (Gatan, Inc., Pleasanton, CA) for 20 seconds in air prior to the application of 3 to 3.5 μL of sample. Grids were blotted and plunge-frozen into liquid ethane with 100% chamber humidity at 5.2 °C using a Vitrobot Mark IV (FEI, Hillsboro, OR) instrument. Eσ^70^ and σP_R_ DNA were on ice for 45 to 100 minutes during grid preparation.

### Cryo-EM data acquisition and processing

Structural biology software was accessed through the SBGrid consortium ^79^. All datasets were collected at the Simons Electron Microscopy Center (SEMC; New York, NY) and recorded using Leginon ^80^. Grids were imaged using a Titan Krios (300 kV accelerating voltage; ThermoFisher Scientific) equipped with a BioQuantum Imaging filter (slit width 20 eV) and either a K3 (tr-Spotiton datasets) or a K2 (5°C dataset) direct electron detector (Gatan, Inc., Pleasanton, CA). All tr-Spotiton grid images were recorded in counting mode with a physical pixel size of 0.844 Å; defocus ranges are listed in Table 1. 5°C grid images were recorded in super-resolution with a physical pixel size of 1.10 Å with a defocus range of -1.0 to -2.3 μm. Dose-fractionated movies were gain-normalized, drift-corrected, summed, and dose-weighted using MotionCor2 ^81^. Data were processed using RELION ^82^ and cryoSPARC (CS) ^83^ (Extended Data Figs. 2-4)

*Eσ*^70^*-σP_R_ tr-Spotiton - 120 ms mixing time.* Datasets 1_120ms_ and 2_120ms_ were independently processed using the pipeline outlined in Extended Data Fig. 2a. After motion correction, the Contrast Transfer Function (CTF) for datasets 1_120ms_ (2,703 micrographs) and 2_120ms_ (2,614 micrographs) was estimated using the Patch CTF module in CS ^83^. CS Blob picker (150-300 Å, local maxima 500) followed by particle extraction (Extract from Micrographs; box size 384 px) yielded an initial set of 593,407 particles (p) (1_120ms_) and of 579,917 p (2_120ms_). Class averages after two rounds of CS 2D classification (N=100 classes) were used to select a subset of particles for reference-free CS Ab-initio Reconstruction (N=3). The ab-initio models (1=RNAP, 2=decoy, 3=decoy) were used to curate all extracted particles using CS Heterogeneous Refinement (N=6, each ab-initio model supplied twice). After two rounds of heterogeneous refinement, 119,486 p (1_120ms_) and 122,054 p (2_120ms_) were refined using CS Non-uniform (NU) refinement ^84^. These particles then underwent one round of Bayesian polishing in RELION ^73^. The polished particles were re-imported into CS where they were NU-refined together (3.0 A, 241,540 p).

To increase the number of particles for later classification strategies, grids from an independent 120 ms tr-Spotiton experiment were imaged and processed (Extended Data Fig. 2b). For dataset 3_120ms_, 21,630 movies (Table 1) were motion-corrected and the CTF was estimated using CS Patch CTF. Micrographs with a CTF fit resolution > 10 were eliminated (2,524). CS Blob picker (150-300 Å, local maxima 500) followed by particle extraction (Extract from Micrographs; box size 256 px) from the remaining micrographs (19,106) yielded 5,731,694 p. Two rounds of CS 2D classification (N=100 classes followed by N=50) were used to select a subset of particles for CS Ab-initio Reconstruction (N=3). The ab-initio models (1=decoy, 2=decoy, 3=RNAP) were used to curate all extracted particles using CS Heterogeneous Refinement (N=6, each ab-initio model supplied twice). After four rounds of heterogeneous refinement, 372,303 p were re-extracted with a boxsize of 384 px, NU refined (nominal resolution of 3.6 Å) and further processed in RELION where they underwent two rounds of Bayesian polishing ^73^. Polished particles were NU refined (nominal resolution of 3.3 A). Polished particles from datasets 1_120ms_, 2_120ms_ and 3_120ms_ were aligned in Class3D (N=1) using the same input volume, 3D auto-refined, and then combined (Join Star) in RELION ^82^ (Extended Data Fig. 4).

*Eσ*^70^*-1P_R_ tr-Spotiton - 500 ms mixing time.* To examine whether the relative populations of intermediates change at longer mixing times, a 500 ms tr-Spotiton dataset was obtained and imaged (Table 1). After motion correction, CTF for dataset 4_500ms_ (14,506 micrographs) was estimated using the CS Patch CTF module ^83^. Micrographs with a CTF fit resolution > 10 were eliminated, leaving 13,511 micrographs. CS Blob picker (150-300 Å, local maxima 400) followed by particle extraction (Extract from Micrographs; box size 256 px) yielded an initial set of 2,787,559 p. Two rounds of CS 2D classification (N=100, N=200 classes) were used to select particles for CS Ab-initio Reconstruction (N=3). The ab-initio models (1=decoy, 2=decoy, 3=RNAP) were used to curate the extracted particles using CS Heterogeneous Refinement (N=6, each ab-initio model supplied twice). After three rounds of heterogeneous refinement, 304,074 p were re-extracted with box size of 384 px, refined using CS Non-uniform (NU) refinement ^84^, and then further processed in RELION with two rounds of Bayesian polishing ^85^. The polished particles were NU refined in CS (3.0 A) (Extended Data Fig. 3).

*3D classification with subtraction.* Polished particles from the 120 ms or the 500 ms datasets were first independently classified using 3D masked classification with subtraction in RELION ^86^. A soft mask encompassing regions that define or bind in the RNAP active site channel was constructed using Chimera ^87^. The “channel” mask included the following: σ^70^_, σ70, σ70_ and σ^70^ (excluding the σ^70^,), the σ^70^_NCR_ βprotrusion and βlobe, the clamp and β’jaw (excluding SI3), and a model of DNA from - 17 to +15 based on PDB 6EE8 ^19^. Classification of subtracted particles did not yield readily interpretable results for the 120 ms datasets. For the 500 ms dataset, 3D classification within the channel mask (N=3, without alignment) after subtraction yielded a class of low-resolution “junk” particles (16%) and two higher resolution classes (84%; Extended Data Fig. 3). The two high-quality classes were distinguished by differences in the cryo-EM density in the downstream channel. Particles from these two classes were combined and used in further processing (Extended Data Figs. 3 and 4).

*3D Variability Analysis (3DVA).* To assess whether discrete states (intermediates) were populated at 120 and 500 ms or whether the conformational landscape at early times is better described as continuous flexibility, the tr-Spotiton datasets were processed using the CS 3DVA algorithm ^21^. Variability components (N=3 components) estimated from the entire consensus 3D maps were dominated by motions in DNA upstream of the -35 hexamer and in the σ^70^ (120 ms or 500 ms). To focus on differences in the active site channel, we first analyzed the 120 and 500 ms datasets independently by locally aligning the particle images in the channel mask, performing 3DVA in the mask, followed by Gaussian mixture model (GMM)-based clustering (N=8). Four discrete classes were detected at each time (detailed below).

For the final processing reported here, the following pipeline was executed. First, we combined the 120 ms and 500 ms particles, reasoning that any conformational differences as a function of time would be robust enough to be sorted during 3DVA classification, and that a larger dataset might amplify low population states. Because the tr-Spotiton sprays do not completely overlap on the grid and because free RNAP may exist in the population in equilibrium with bound DNA, we eliminated free RNAP particles by constructing a soft “DNA” mask around the promoter DNA from ∼-40 to -10, the proximal σCTD, and σ^70^ . After signal subtraction using the DNA mask, the combined particle stack underwent masked 3D classification (RELION, N=6 classes, without alignment) followed by NU refinement (CS), yielding a low resolution “junk” class (6%), one free RNAP class (13%), and four DNA bound classes (80%; Extended Data Fig. 4).

DNA-bound particles (635,069 p) were combined, NU refined, and then locally aligned in CS using the channel mask. The resulting particle stack was then subjected to 3DVA (N=3, filter resolution=6 Å). GMM-based clustering of the variability component corresponding to clamp opening and closing movements (N=8; see Main Text, Supplemental Video 2) revealed four distinct conformational states, distinguished primarily by different degrees of clamp closure, extents of resolution of the single stranded nt-strand and changes in the cryo-EM density in the downstream channel. Particles corresponding to each class were combined and NU refined with per-particle defocus refinement and Global CTF refinement enabled (Extended Data Fig. 4).

*Eσ*^70^*-1P_R_ tr-Spotiton - 500 ms using fluorinated Fos-Choline-8 (FC8F).* Because of its effectiveness in preventing preferred particle orientations, CHAPSO has been used in cryo-EM studies, particularly those of *Eco* RNAP ^26^. However, CHAPSO binds multiple sites on *Eco* RNAP ^26^, including the σ-finger (σ^70^) in the active site cleft ^7^. To examine whether CHAPSO influenced the nature of the intermediates captured in tr-Spotiton, we (A.U.M.) found FC8F (1.5 mM) acts as effectively as CHAPSO to mitigate particle orientation bias but does not interact with Eσ^70^. Cryo-EM grids obtained from a 500 ms tr-Spotiton experiment where 1.5 mM FC8F replaced CHAPSO were imaged and processed using the same pipeline used for the 500 ms data in CHAPSO (Extended Data Fig. 8). The same intermediates were found, indicating that the results are independent of CHAPSO.

*Eσ*^70^*-1P_R_ complexes populated at 5°C.* Cryo-EM grids prepared by vitrifying complexes populated at equilibrium at low temperature were imaged and processed using the pipeline shown in Extended Data Fig. 9.

Local resolution-filtered cryo-EM maps were generated using blocres and blocfilt from the Bsoft package ^75^ with the following parameters: box size 20, sampling using the physical pixel size (Extended Data Figs. 6, 7, 9; Table 1). Directional 3DFSCs were calculated using 3DFSC ^88^ (Extended Data Figs. 6, 7, 9).

### Model building and refinement

To build initial models of the protein-DNA components in the tr-Spotiton cryo-EM density maps, Eσ^70^ bound to 1P_R_ from PDB 7MKD ^7^ was manually fit into the density map using Chimera ^87^ and real-space refined using PHENIX ^89^. The Eσ^70^-*rpsT* P2 promoter closed complex (RPc) (PDB 6PSQ) ^10^ served as the initial model for the RPc intermediate populated at 5°C at 1P_R_. Real-space refinement, rigid body refinement with sixteen manually-defined mobile domains was followed by all-atom and B-factor refinement with Ramachandran and secondary structure restraints. Models were inspected and modified in Coot ^90^. Alignment shown in Fig. 1b was done using conserved domains of RNAP that exhibit minimal conformational changes in the transcription cycle (the RNAP structural core) ^91^. Statistical analyses for model refinement and validation were generated using MolProbity ^92^ and PHENIX ^89^.

### Disulfide crosslinking and abortive transcription assays

*Purification of σ*^70^ *region 1.1 cysteine pair variants.* A cysteine-less variant of *Eco* σ^70^ (C132S, C291S, C295S) and variants containing cysteine pair mutations at I35C-S89C, Q8C-P32C, or Y21C-Q54C were expressed with a His_10_-SUMO N-terminal tag from a pET28-based vector in *Eco* BL21(DE3). 2 L shaking cultures inoculated from overnight cultures of freshly transformed cells were grown at 37°C to OD600 of 0.7 to 1.0, at which point the temperature was reduced to 25°C. Expression was induced after 1 h with a final concentration of 0.5 mM IPTG and continued overnight at 25°C. Cells were harvested by centrifugation and resuspended in about 20 ml/L culture of buffer A [20 mM Tris-HCl pH 8.0/RT, 1 M NaCl, 5% glycerol (v/v), 2 mM DTT] supplemented with 5 mM imidazole, 1 mM phenylmethylsulfonyl flouride (PMSF, Sigma-Aldrich) and 1x protease inhibitor cocktail (c0mplete EDTA-free, Roche). Cells were lysed using a continuous-flow homogenizer (Avestin) at 4°C. After centrifugation of the lysate at 15,000 rpm (JA-18 rotor; ∼33,000 x g) for 30 min at 4°C, the soluble fraction was recovered, filtered through 5 μm PVDF membrane (EMD Millipore), and applied to a 5 ml HiTrap IMAC Sepharose FF column (Cytiva) charged with Ni^2+^ ions and equilibrated in buffer B [20 mM Tris-HCl pH 8.0/RT, 1 M NaCl, 5% glycerol (v/v), 2 mM DTT, 500 mM imidazole]. Subsequently, unbound material was eluted from the column by wash steps using buffer A and B of 5 CV 5% B, and 10 CV 10% B. Protein was eluted with 70% B (with 350 mM imidazole) and fractions were analyzed for protein content by SDS-PAGE and Coomassie staining. Protein-containing fractions were pooled and supplemented with 0.5 mM EDTA and 1 mM DTT. To remove the N-terminal tag, His-tagged Ulp1 protease was added at 1:30 to 1:50 molar ratio and the sample was dialyzed in a SpectraPor 12-14 kDa membrane against 2 L buffer A supplemented with 0.1 mM EDTA overnight at 4°C. Ulp1 and uncleaved protein were removed by passing the sample over a 5 ml HiTrap IMAC Sepharose FF column (Cytiva) charged with Ni^2+^ ions. The flow through was concentrated using centrifugal filters with a 30 kDa cutoff membrane (Amicon 30K, EMD Millipore) and applied to a Superdex 200 prep grade 26/600 column (Cytiva) using buffer C [20 mM HEPES-NaOH pH 8.0/RT, 500 mM NaCl, 5% glycerol (v/v), 5 mM DTT, 0.1 mM EDTA]. Peak fractions were collected and pooled according to protein content. Variants were concentrated to 100-250 μM protein using centrifugal filters (Amicon 30K) and the final sample was mixed 1:1 with buffer C containing 50% glycerol (v/v) before freezing aliquots in liquid nitrogen. Frozen protein samples were stored at −80°C until use.

In vitro *abortive transcription assay for σ*^70^ *cysteine pair variants.* Cysteine pair variants and the cysteine-less variant of *Eco* σ^70^ were buffer exchanged to buffer R (40 mM Tris-HCl pH 8/RT, 120 mM KCl, 10 mM MgCl_2_, 20 ug/ml BSA) using Zeba 5K desalting spin columns (ThermoFisher Scientific). The variants were subjected to oxidation by mixing σ^70^ variant at 2.5 μM final concentration with hydrogen peroxide at 5 mM final concentration in buffer R. Reactions (typically 10-20 μl) were incubated for 10 min at 37°C. After the incubation, catalase (1 U/ul) was added to about 0.17 U/ul final concentration to neutralize the hydrogen peroxide and the reactions were incubated at 25°C for at least 5 min. Crosslinking efficiency was assessed by SDS-PAGE using 10% AA/BAA (37.5:1) Bis-Tris gels and Coomassie staining (typically >90%).

Abortive transcription reactions were carried out on a 1P_R_ promoter fragment (-60 to +30) ^7^. First, Eσ^70^ was formed by mixing *Eco* core RNAP with σ^7-^ variant (and DTT for the reaction under reducing conditions) in buffer R at 37°C for 10 min. DNA was added to start formation of open complexes. Final concentrations of each component were 40 nM core RNAP, 100 nM σ^7-^ variant, 10 nM DNA, and 1 mM DTT for the reactions under reducing conditions (DTT was replaced with buffer R for oxidizing conditions). After incubating the reaction at 25°C for 90 seconds, the complexes were challenged with heparin (50 ug/ml final) for 1-2 min (depending on the handling time). NTP mix (250 uM ApU, 50 uM GTP, 130 nCi/μl σ-^32^P-GTP, 50ug/mL heparin) was added to start abortive transcription. Reactions were incubated for 5 min at 25°C before mixing with an equal volume of 2x STOP buffer [0.5x TBE, 8 M urea, 30 mM EDTA, 0.05 % bromophenol blue (w/v), 0.05 % xylene cyanol (w/v]) and heating to 95°C to stop the reaction. Products were analyzed by loading 4.5 μl sample on a 23% TBE-urea gel run in 1x TBE for 1:30 h at 1000 V. Gels were exposed to a storage phosphor screen overnight at 4°C. Imaging was performed on a Typhoon imager (Cytiva).

### Data Availability

All unique/stable reagents generated in this study are available without restriction from the Lead Contact, Seth A. Darst. The cryo-EM density maps and atomic coordinates have been deposited in the EMDataBank and Protein Data Bank as follows: RPc_5°C_ (EMD-41456, 8TOM), I1a (EMD-41433, 8TO1), I1b (EMD-41439, 8TO8), I1c (EMD-41448, 8TOE), I1d (EMD-41437, 8TO6).

## EXTENDED DATA

**Extended Data Fig. 1.**
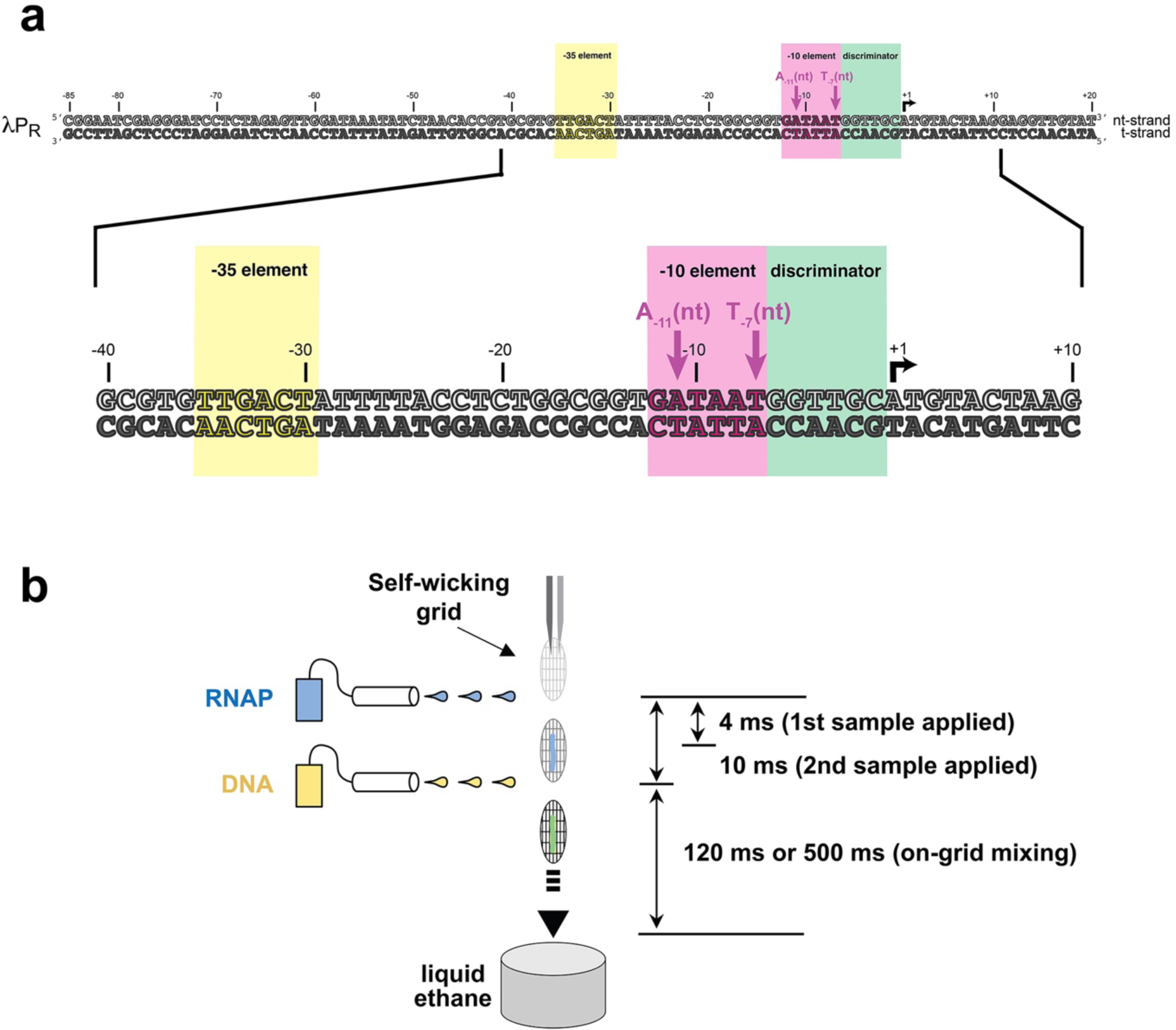
1P_R_ promoter fragment and tr-Spotiton. **a.** 1P_R_ promoter DNA construct (-85 to +20) used for cryo-EM studies. The sequence from -40 to +10 is magnified below. **b.** Schematic diagram illustrating the principle of the tr-Spotiton device. For more details see ref. ^20^.

**Extended Data Fig. 2.**
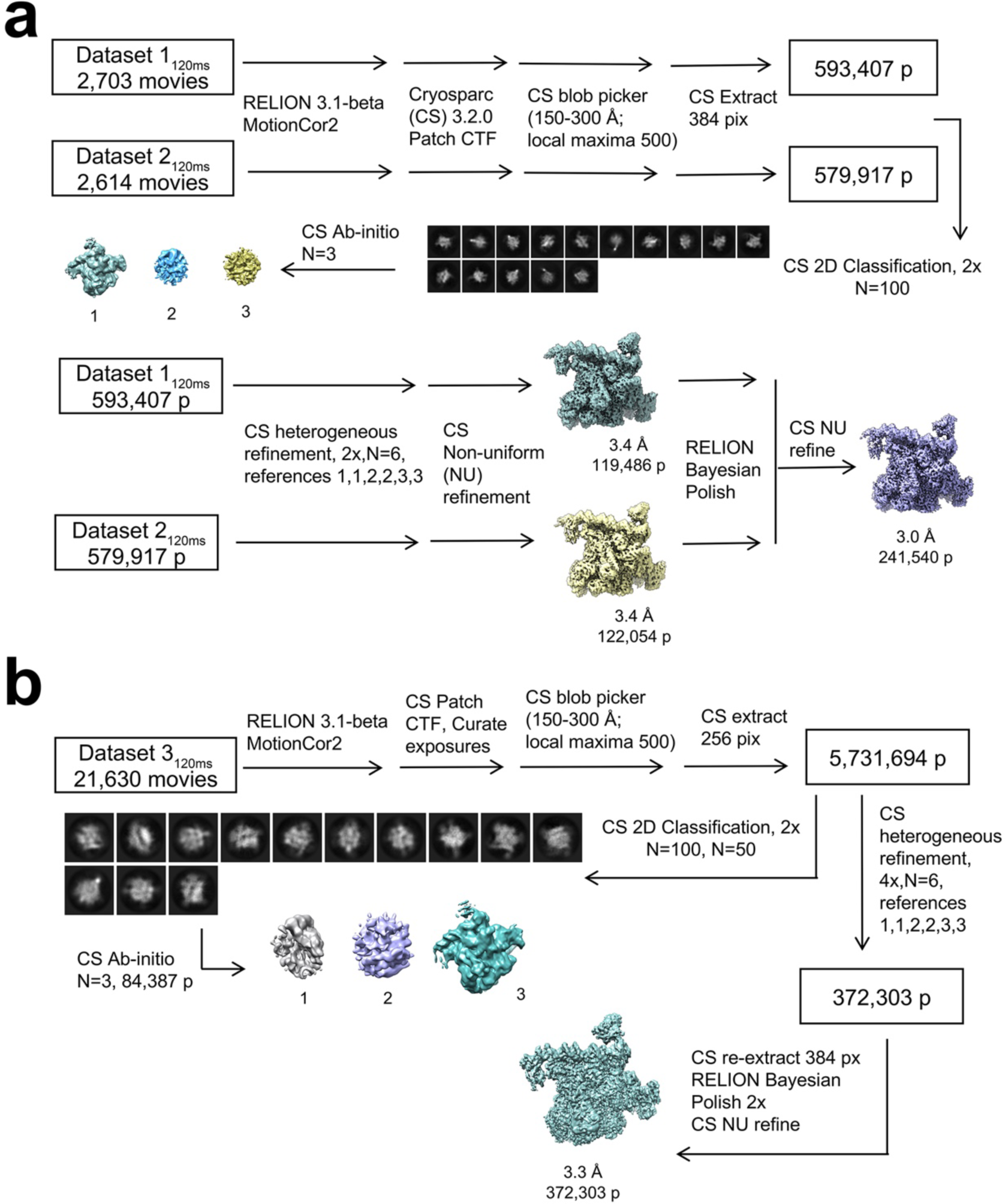
Cryo-EM processing pipeline for 120 ms datasets. Cryo-EM processing pipelines for *Eco* RNAP mixed with 1P_R_ DNA using tr-Spotiton (t = 120 ms, 8 mM CHAPSO) **a.** Datasets 1_120ms_ and 2_120ms_. **b.** Dataset 3_120ms_.

**Extended Data Fig. 3.**
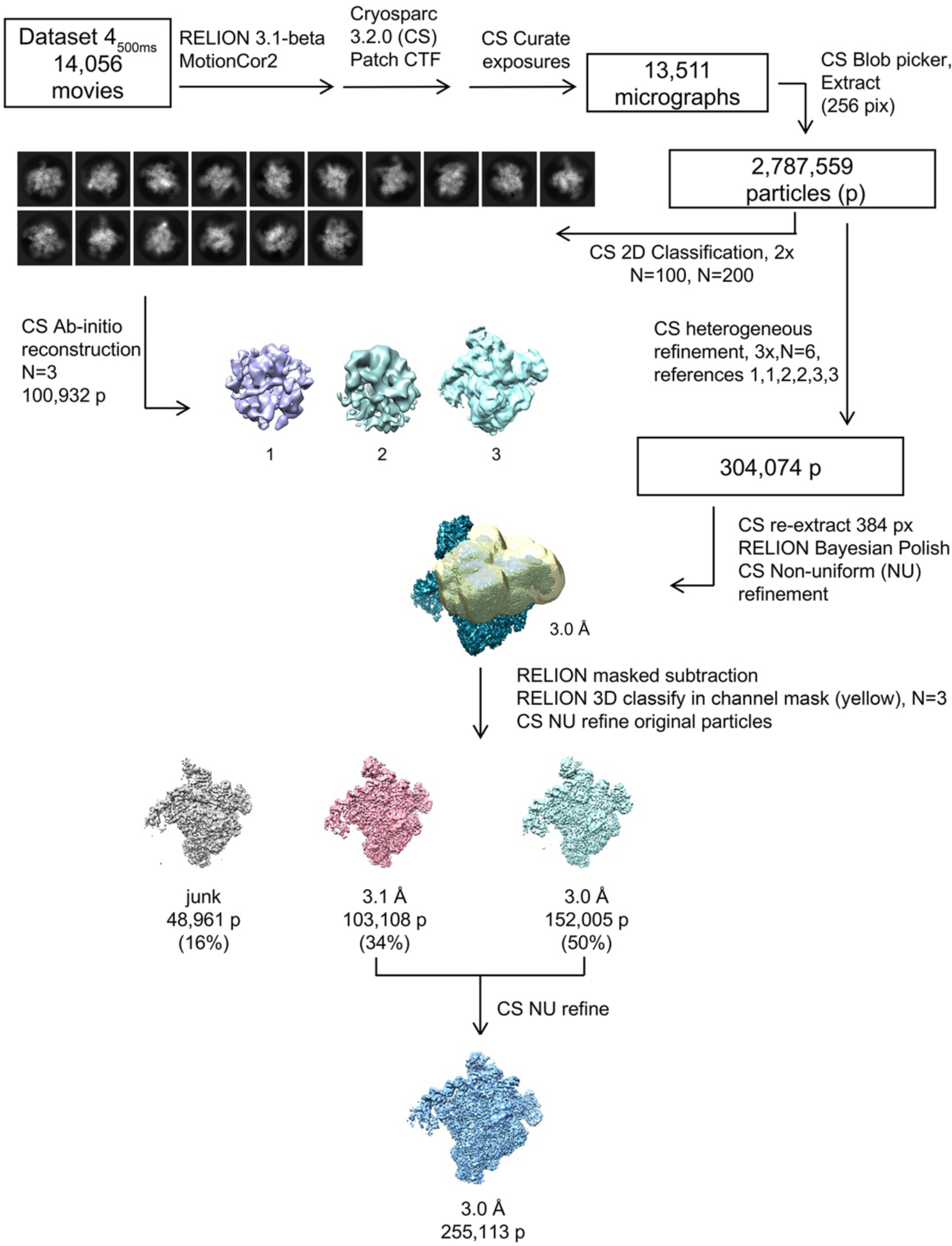
Cryo-EM processing pipeline for dataset 4_400ms_. Cryo-EM processing pipeline for *Eco* RNAP mixed with 1P_R_ DNA using tr-Spotiton (t = 500 ms, 8 mM CHAPSO, dataset 4).

**Extended Data Fig. 4.**
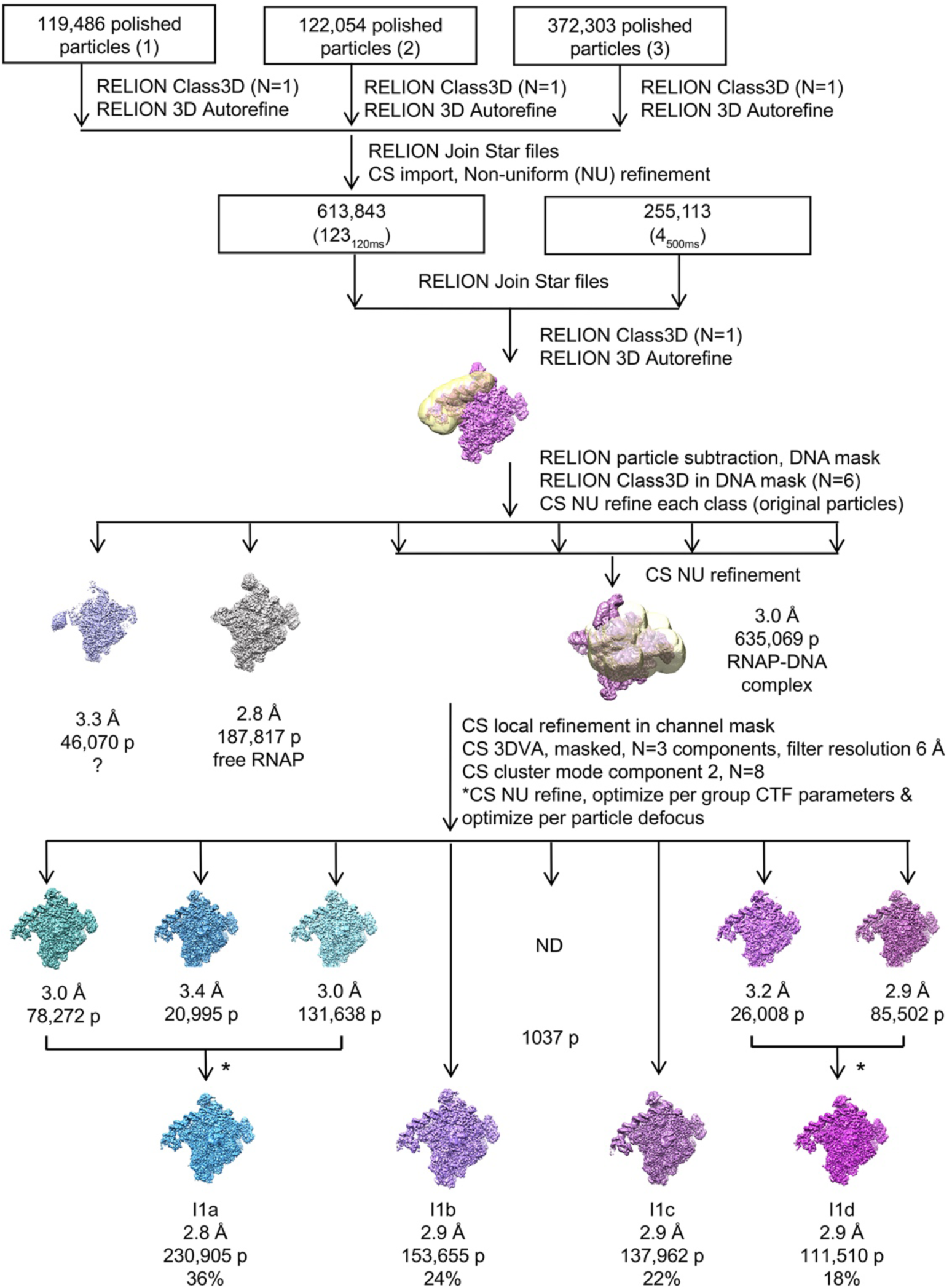
Cryo-EM processing pipeline for combined datasets. Cryo-EM processing pipeline for combining polished particles from Spotiton datasets 1 - 3 (t = 120 ms; see Extended Data Fig. 2) and dataset 4 (t = 500 ms; see Extended Data Fig. 3).

**Extended Data Fig. 5.**
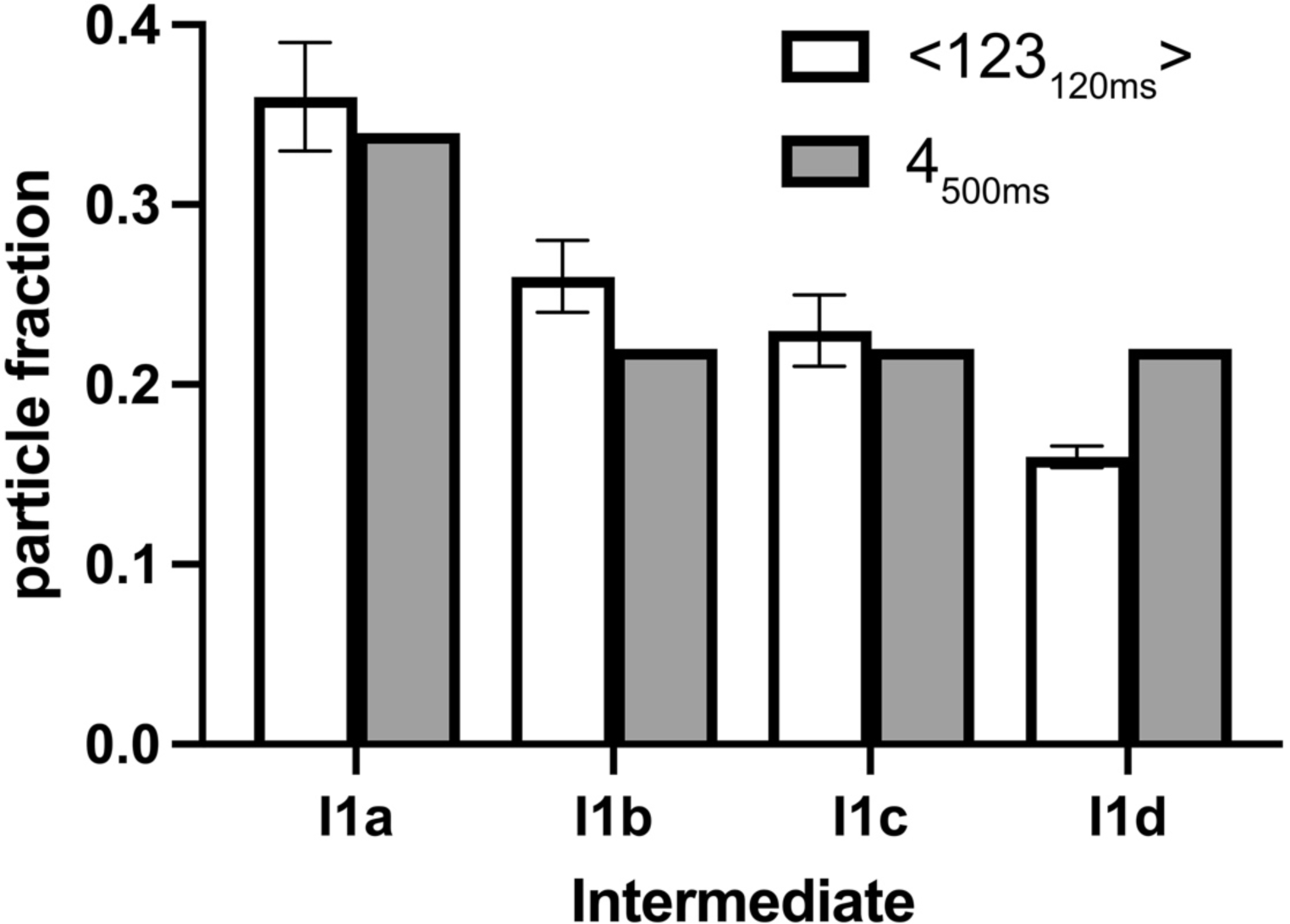
Comparison of particle population distributions for 120 ms vs. 500 ms datasets. Histogram plots showing the fraction of particles that contribute to each intermedate. The open bars show the mean particle fraction for the three 120 ms datasets (<123_120ms_>); the error bars denote the standard deviation for n=3. The gray bars denote the particle fraction for the single 500 ms dataset (4_500ms_).

**Extended Data Fig. 6.**
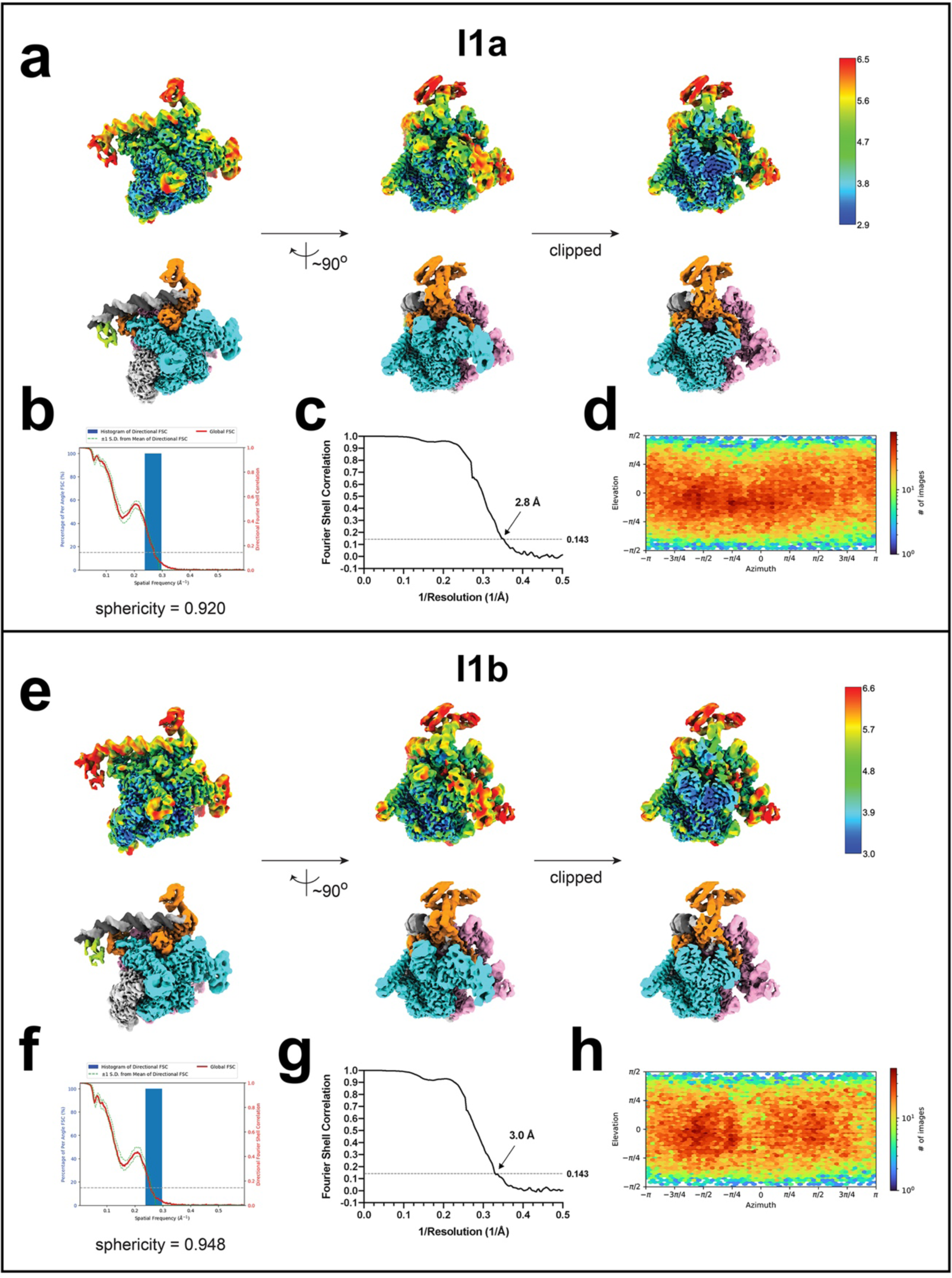
Cryo-EM of I1a and I1b. **a.-d.** Cryo-EM of I1a. **a.** Three views of the combined nominal 2.8 Å resolution cryo-EM map, filtered by local resolution ^75^. The bottom row is colored according to the subunit (σI, σII, light grey; σCTD, limon; β, cyan; β’, pink; σ^70^, orange; DNA t-strand, dark grey; DNA nt-strand, grey). The top row shows the same views but colored by local resolution ^75^. The right view is a cross-section through the middle view. **b.** Directional 3D FSC, determined with 3DFSC ^88^. **c.** Gold-standard FSC plot ^93^, calculated by comparing two half maps. The dotted line represents the 0.143 FSC cutoff. **d.** Angular distribution plot, calculated in cryoSPARC. Scale depicts number of particles assigned to a specific angular bin. **e.-h.** Cryo-EM of I1b. **e.** Three views of the combined nominal 3.0 Å resolution cryo-EM map, filtered by local resolution ^75^. The bottom row is colored according to the subunit (σI, σII, light grey; σCTD, limon; β, cyan; β’, pink; σ^70^, orange; DNA t-strand, dark grey; DNA nt-strand, grey). The top row shows the same views but colored by local resolution ^75^. The right view is a cross-section through the middle view. **f.** Directional 3D FSC, determined with 3DFSC ^88^. **g.** Gold-standard FSC plot ^93^, calculated by comparing two half maps. The dotted line represents the 0.143 FSC cutoff. **h.** Angular distribution plot, calculated in cryoSPARC. Scale depicts number of particles assigned to a specific angular bin.

**Extended Data Fig. 7.**
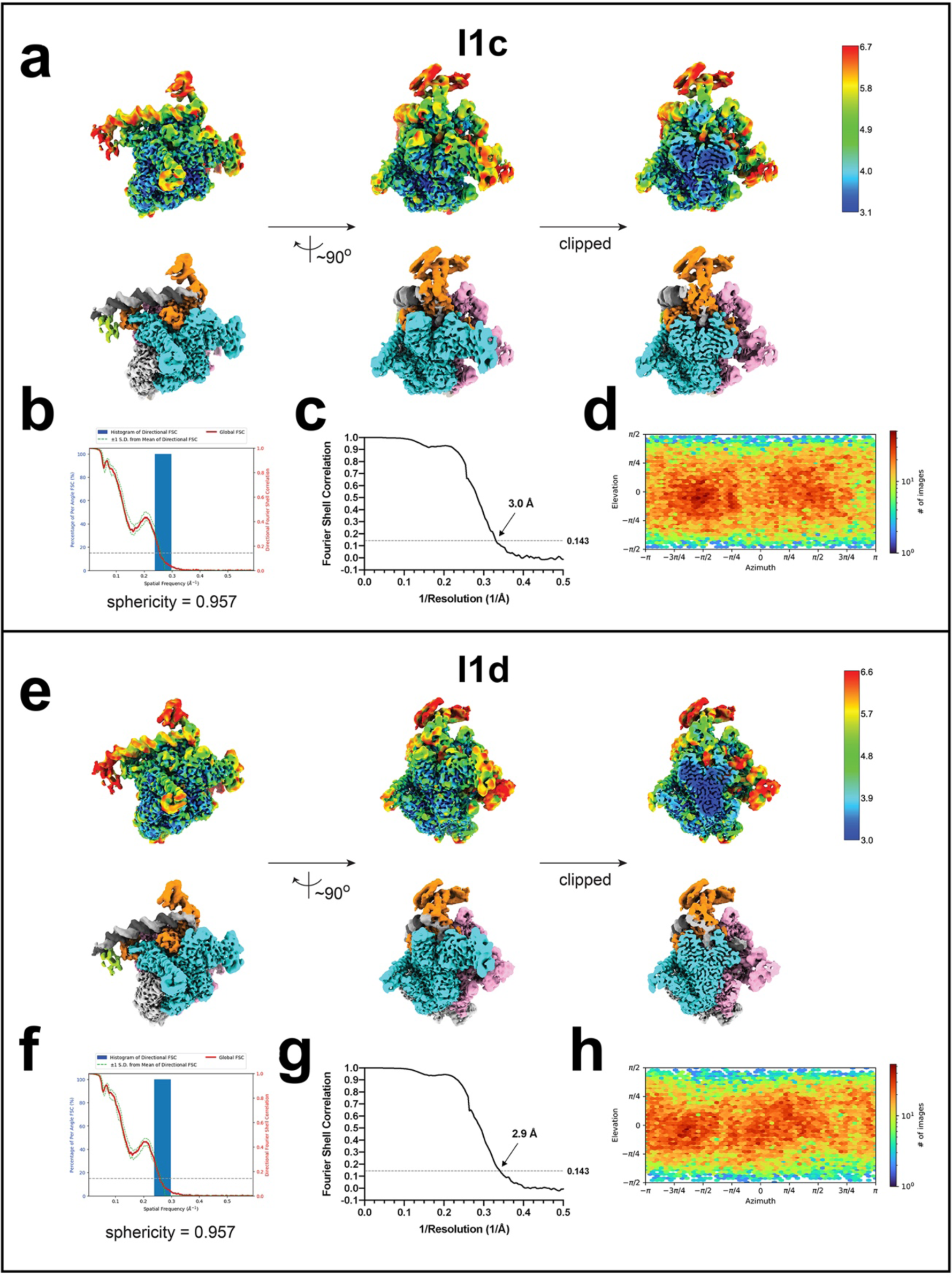
Cryo-EM of I1c and I1d. **a.-d.** Cryo-EM of I1c. **a.** Three views of the combined nominal 3.0 Å resolution cryo-EM map, filtered by local resolution ^75^. The bottom row is colored according to the subunit (σI, σII, light grey; σCTD, limon; β, cyan; β’, pink; σ^70^, orange; DNA t-strand, dark grey; DNA nt-strand, grey). The top row shows the same views but colored by local resolution ^75^. The right view is a cross-section through the middle view. **b.** Directional 3D FSC, determined with 3DFSC ^88^. **c.** Gold-standard FSC plot ^93^, calculated by comparing two half maps. The dotted line represents the 0.143 FSC cutoff. **d.** Angular distribution plot, calculated in cryoSPARC. Scale depicts number of particles assigned to a specific angular bin. **e.-h.** Cryo-EM of I1d. **e.** Three views of the combined nominal 2.9 Å resolution cryo-EM map, filtered by local resolution ^75^. The bottom row is colored according to the subunit (σI, σII, light grey; σCTD, limon; β, cyan; β’, pink; σ^70^, orange; DNA t-strand, dark grey; DNA nt-strand, grey). The top row shows the same views but colored by local resolution ^75^. The right view is a cross-section through the middle view. **f.** Directional 3D FSC, determined with 3DFSC ^88^. **g.** Gold-standard FSC plot ^93^, calculated by comparing two half maps. The dotted line represents the 0.143 FSC cutoff. **h.** Angular distribution plot, calculated in cryoSPARC. Scale depicts number of particles assigned to a specific angular bin.

**Extended Data Fig. 8.**
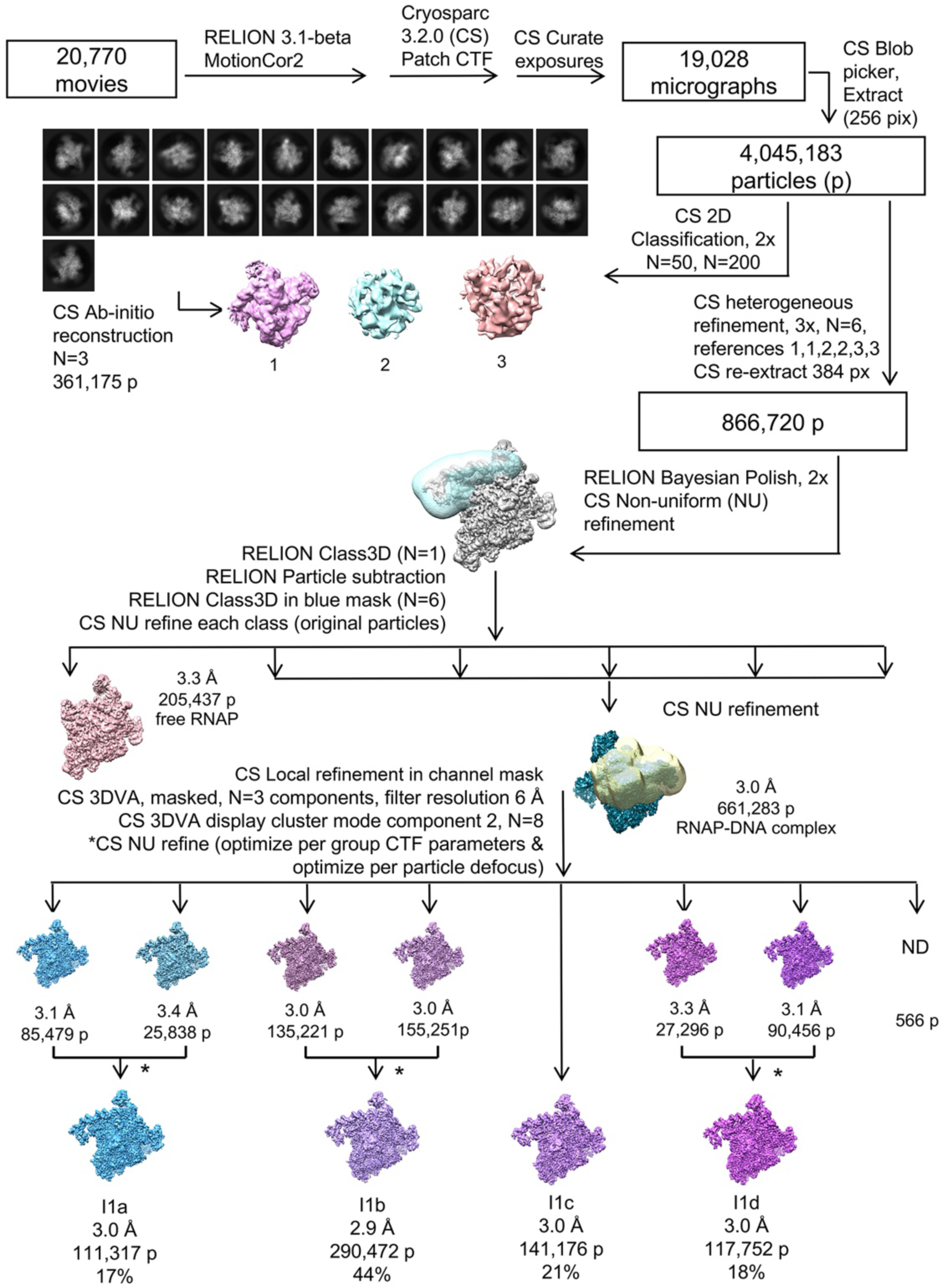
Cryo-EM processing pipeline for dataset 5_500ms,FC8F_. Cryo-EM processing pipeline for *Eco* RNAP mixed with 1P_R_ DNA using tr-Spotiton (t = 500 ms, 1.5 mM FC8F).

**Extended Data Fig. 9.**
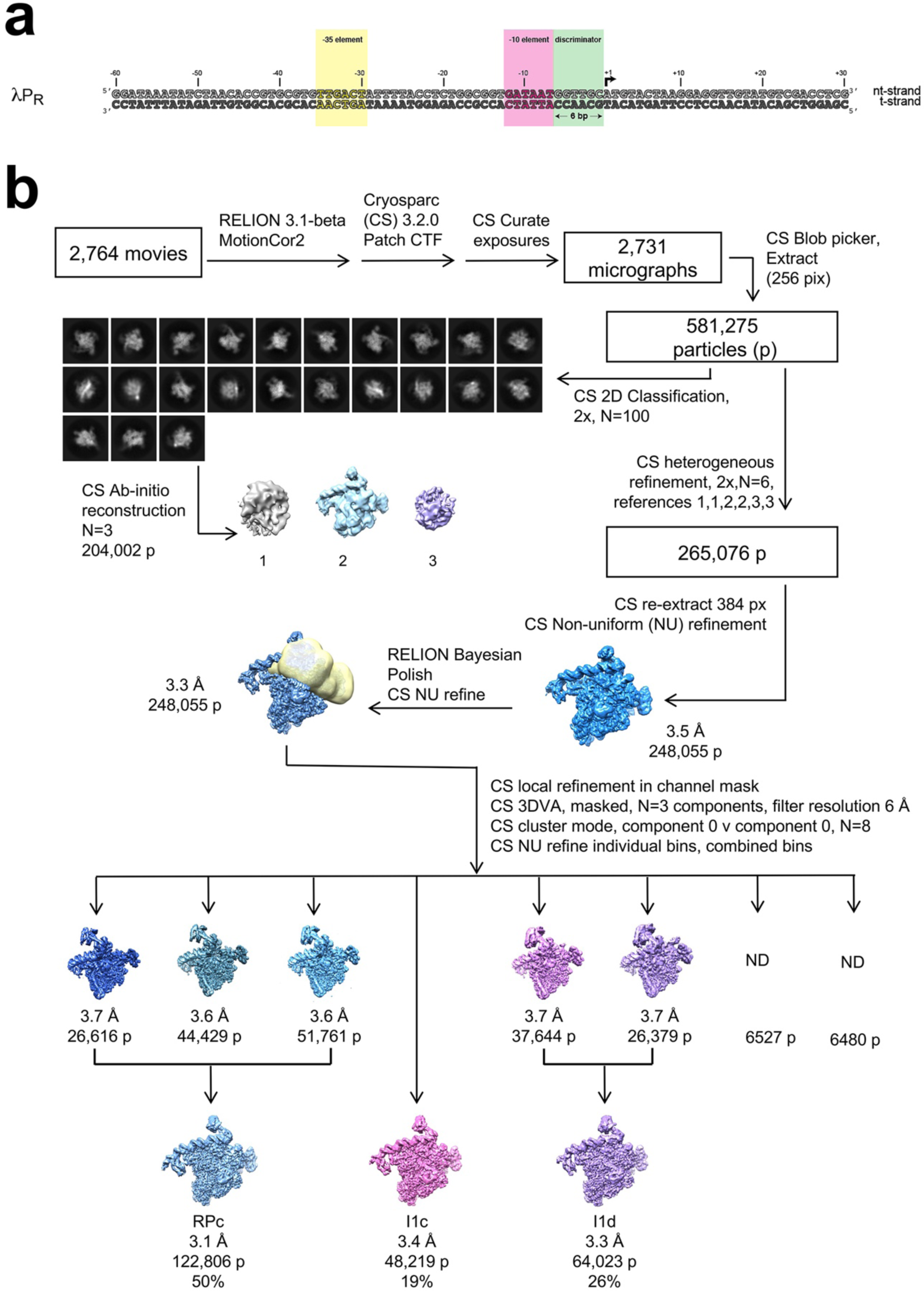
Cryo-EM processing pipeline for 5°C dataset. **a.** 1P_R_ promoter DNA construct used for 5°C cryo-EM studies. **b.** Cryo-EM processing pipeline for *Eco* RNAP and 1P_R_ DNA (-60 to +30) mixed manually and allowed to come to equilibrium at 5°C (See Methods).

**Extended Data Fig. 10.**
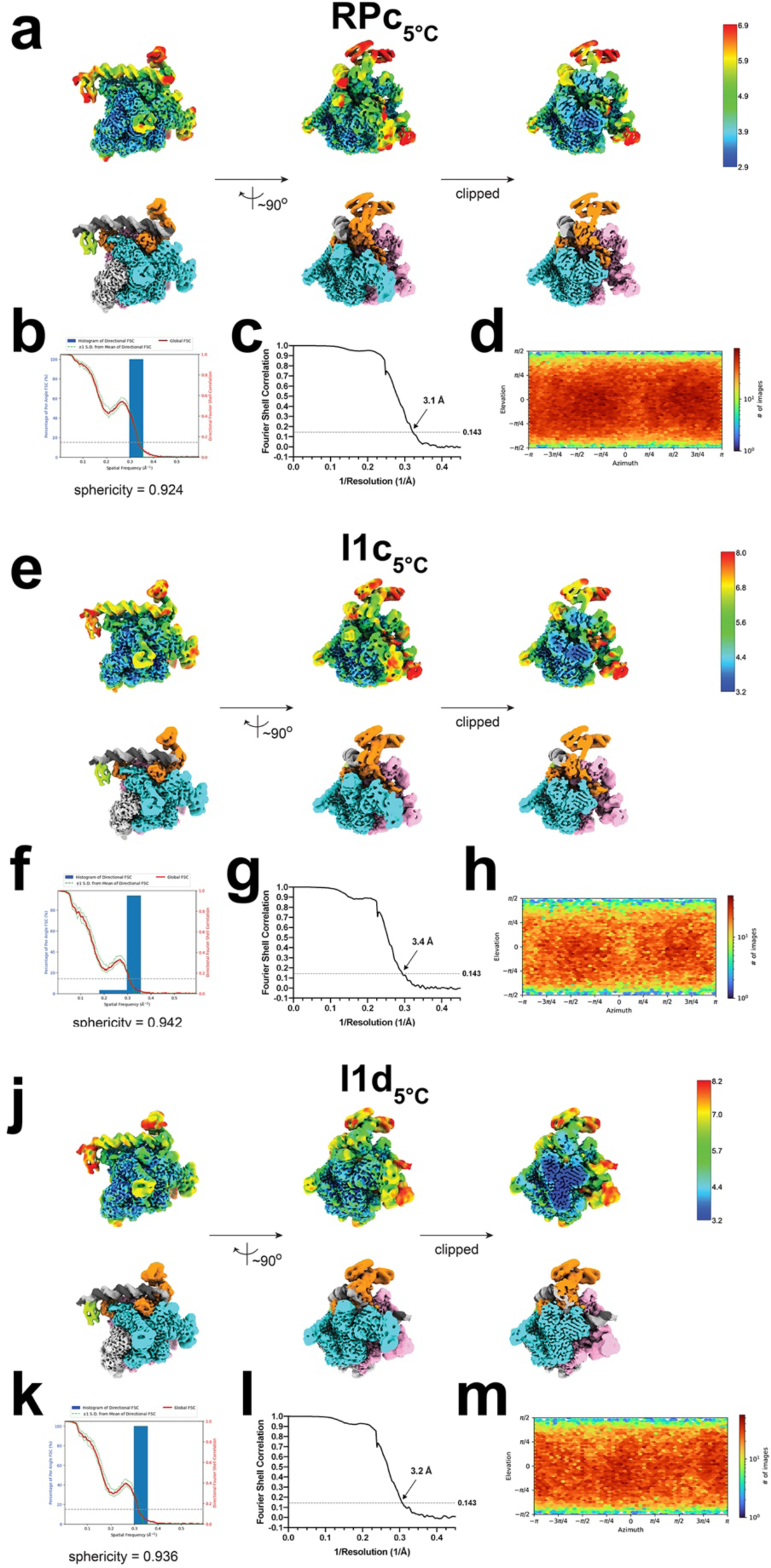
Cryo-EM of RPc_5°C_, I1c_5°C_, and I1d_5°C_. **a.-d.** Cryo-EM of RPc_5°C_. **a.** Three views of the combined nominal 3.1 Å resolution cryo-EM map, filtered by local resolution ^75^. The bottom row is colored according to the subunit (σI, σII, light grey; σCTD, limon; β, cyan; β’, pink; σ^70^, orange; DNA t-strand, dark grey; DNA nt-strand, grey). The top row shows the same views but colored by local resolution ^75^. The right view is a cross-section through the middle view. **b.** Directional 3D FSC, determined with 3DFSC ^88^. **c.** Gold-standard FSC plot ^93^, calculated by comparing two half maps. The dotted line represents the 0.143 FSC cutoff. **d.** Angular distribution plot, calculated in cryoSPARC. Scale depicts number of particles assigned to a specific angular bin. **e.-h.** CryoEM of I1c_5°C_. **e.** Three views of the combined nominal 3.4 Å resolution cryo-EM map, filtered by local resolution ^75^. The bottom row is colored according to the subunit (σI, σII, light grey; σCTD, limon; β, cyan; β’, pink; σ^70^, orange; DNA t-strand, dark grey; DNA nt-strand, grey). The top row shows the same views but colored by local resolution ^75^. The right view is a cross-section through the middle view. **f.** Directional 3D FSC, determined with 3DFSC ^88^. **g.** Gold-standard FSC plot ^93^, calculated by comparing two half maps. The dotted line represents the 0.143 FSC cutoff. **h.** Angular distribution plot, calculated in cryoSPARC. Scale depicts number of particles assigned to a specific angular bin. **j.-m.** CryoEM of I1d_5°C_. **j.** Three views of the combined nominal 3.2 Å resolution cryo-EM map, filtered by local resolution ^75^. The bottom row is colored according to the subunit (σI, σII, light grey; σCTD, limon; β, cyan; β’, pink; σ^70^, orange; DNA t-strand, dark grey; DNA nt-strand, grey). The top row shows the same views but colored by local resolution ^75^. The right view is a cross-section through the middle view. **k.** Directional 3D FSC, determined with 3DFSC ^88^. **l.** Gold-standard FSC plot ^93^, calculated by comparing two half maps. The dotted line represents the 0.143 FSC cutoff. **m.** Angular distribution plot, calculated in cryoSPARC. Scale depicts number of particles assigned to a specific angular bin.

**Extended Data Fig. 11.**
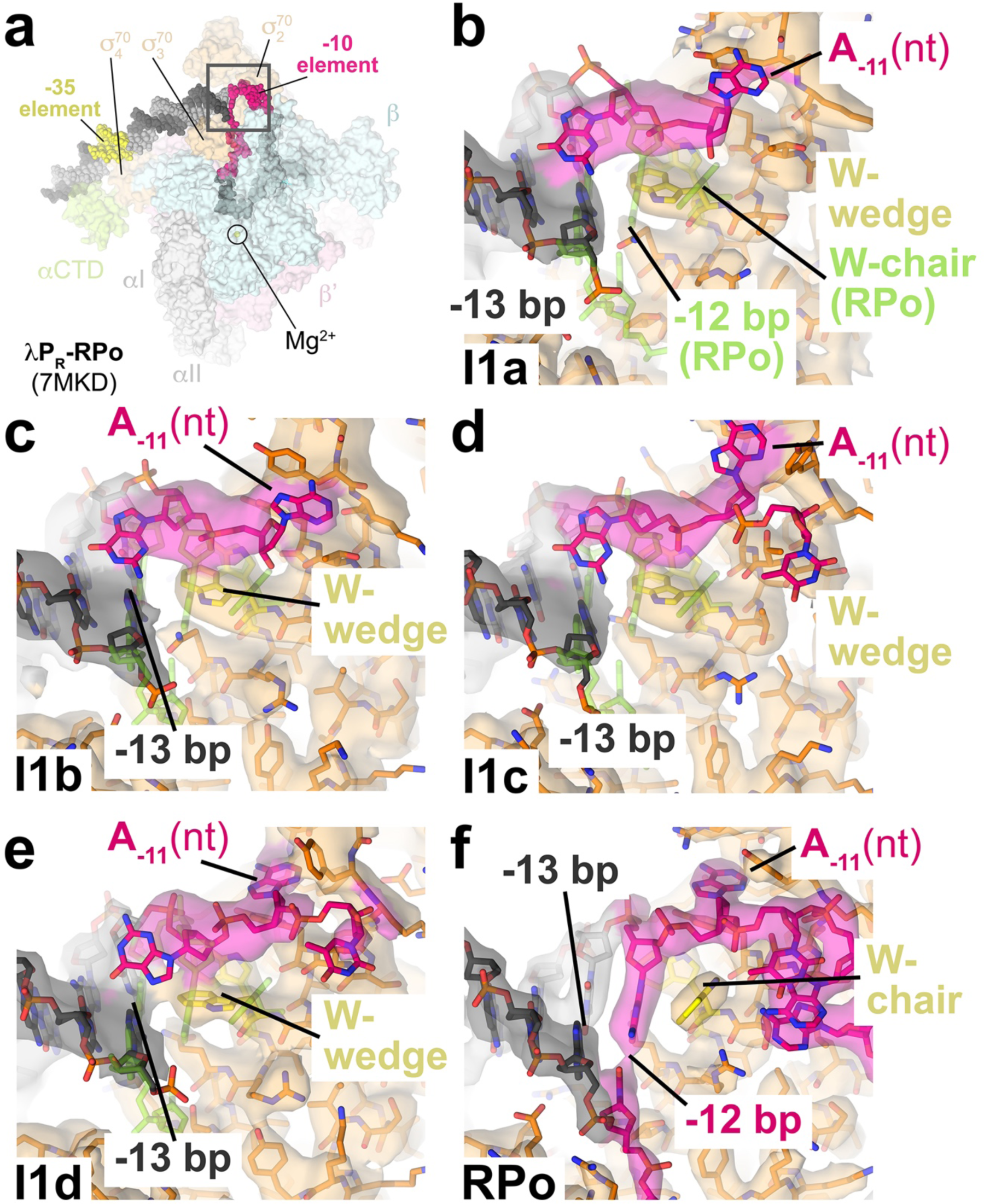
The σ^70^ W-dyad and the -12 bp in 1P_R_ intermediates. **a.** Top view of 1P_R_-RPo (7MKD) ^7^. Eσ^70^ is shown as a transparent molecular surface. The DNA is shown as atomic spheres, color-coded as in Fig. 2a. **b.-f.** The boxed region in (**a**) is magnified, showing the region of the σ^70^ W-dyad and the -13 to -11 positions of the promoter. σ^70^ and DNA (color-coded as in Fig. 3) are shown in stick format; σ^70^ carbon atoms are colored orange but the W-dyad is highlighted in yellow. Transparent cryo-EM density (local-resolution filtered ^75^) is superimposed. For reference, the positions of key RPo elements are shown in stick format and colored chartreuse (W-dyad in chair conformation and the -12 bp). For I1a, I1b, I1c, and I1d (**b.-e.**), the W-dyad is in the edge-on (wedge) conformation and the -12 bp is opened. Only in RPo is the W-dyad in the chair conformation and the -12 bp re-paired.

**Extended Data Fig. 12.**
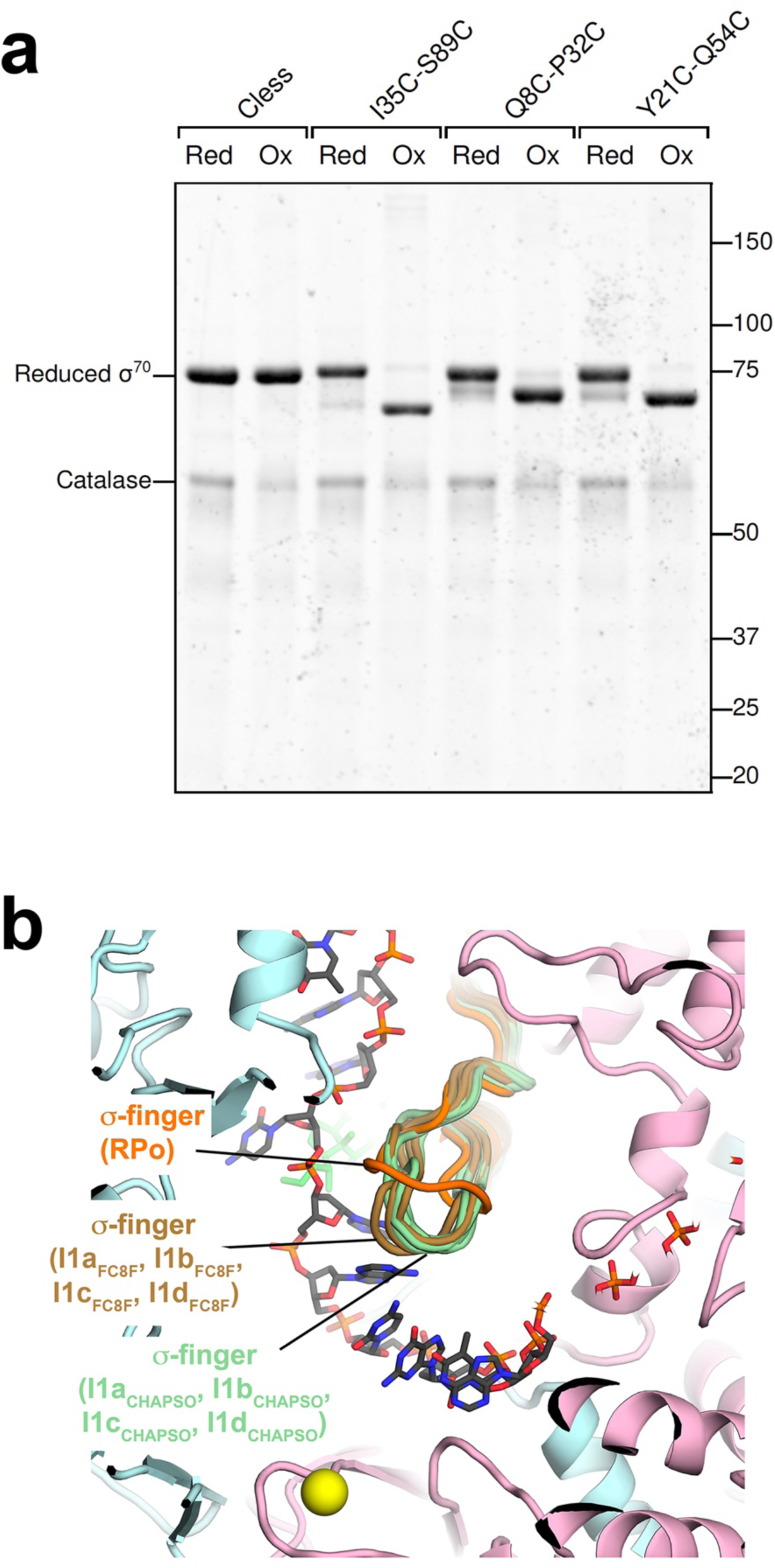
σ^70^_1.1_ disulfide crosslinking, and a conformational change in the σ-finger. **a.** σ^70^ derivatives were analyzed by 10% SDS-polyacrylamide gel electrophoresis and visualized with Coomassie stain. Each σ^70^ derivative was analyzed under reducing (preventing formation of any disulfide bonds) or oxidizing (promoting the formation of disulfide bonds) conditions. Each Cys-pair mutant shows higher mobility under oxidizing conditions indicating formation of the relevant disulfide bond. Moreover, the difference in mobility between the reduced and oxidized condition correlates with the number of residues separating the two engineered Cys substitutions (I35C-S89C, 55 residues; Q8C-P32C, 25 residues; Y21C-Q54C, 34 residues). **b**. The 1P_R_-RPo structure (7MKD) ^7^ in the active-site region is shown; The RNAP is shown as a backbone cartoon (β, light cyan; β’, light pink; σ^70^, orange); t-strand DNA is shown in stick format (carbon atoms dark grey); the RNAP active-site Mg^2+^ is shown as a yellow sphere. The structures of I1a, I1b, I1c, and I1d from the CHAPSO (light green) and FC8F (brown) datasets were superimposed by the RNAP structural core and shown is the σ-finger from each.

