## Supplemental information for "Early intermediates in bacterial RNA polymerase promoter melting visualized by time-resolved cryo-electron microscopy"

The authors declare no conflict of interest

[4] These authors contributed equally.

[7] Current address: Memorial Sloan Kettering Cancer Center, Sloan Kettering Institute, New York, NY 10065 USA.

[8] Current address: Department of Cell Biology, New York University School of Medicine, New York, NY 10016 USA.

[9] Current address: National Institute of Environmental Health Sciences, Durham, NC 27709 USA.

[10] Current address: Chan Zuckerberg Imaging Institute,

Supplemental information includes 2 tables and 2 videos.

**Supplementary Table 1.**

**a. Fraction of particles from combined dataset ($123_{150ms}4_{500ms}$) distributed into each I1 intermediate.**

| Dataset | $t_{mix}$ | fraction of particles | | | |
|---|---|---|---|---|---|
| | | I1a | I1b | I1c | I1d |
| 1 | 150 ms | 0.34 | 0.27 | 0.24 | 0.15 |
| 2 | 150 ms | 0.35 | 0.26 | 0.23 | 0.16 |
| 3 | 150 ms | 0.40 | 0.24 | 0.21 | 0.16 |
| 123 | 150 ms | 0.36 ± 0.03 | 0.26 ± 0.02 | 0.23 ± 0.02 | 0.157 ± 0.006 |
| 4 | 500 ms | 0.34 | 0.22 | 0.22 | 0.22 |

**b. Jensen-Shannon distance analysis of particle distributions.**

| | $1_{150ms}$ | $2_{150ms}$ | $3_{150ms}$ | $4_{500ms}$ |
|---|---|---|---|---|
| $1_{150\ ms}$ | | 0.018 | 0.057 | 0.083 |
| $2_{150\ ms}$ | | | 0.043 | 0.069 |
| $3_{150\ ms}$ | | | | 0.075 |
| $4_{500\ ms}$ | | | | |

$<1_{150ms}|2_{150ms}, 1_{150ms}|3_{150ms}, 2_{150ms}|3_{150ms}> = 0.039 \pm 0.020$

$<1_{150ms}|4_{500ms}, 2_{150ms}|4_{500ms}, 3_{150ms}|4_{500ms} > = 0.076 \pm 0.007$

**Supplementary Table 2. Characteristics of $\lambda P_R$ RPo formation intermediates.**

| | | Intermediate | | | | | |
|---|---|---|---|---|---|---|---|
| | | RPc | I1a | I1b | I1c | I1d | RPo |
| $A_{-11}$(nt) | flipped | No | Yes | Yes | Yes | Yes | Yes |
| | captured | No | No | No | No | Yes | Yes |
| $T_{-7}$(nt) | captured | No | No | Yes | Yes | Yes | Yes |
| downstream channel | | $\sigma^{70}_{1.1}$ | $\sigma^{70}_{1.1}$ | $\sigma^{70}_{1.1}{}^{*}$ | $\sigma^{70}_{1.1}{}^{*}$ | DNA | DNA |
| extent of transcription bubble | | no bubble | ≥ 2 bp (-12 to -11) | ≥ 9 bp (-12 to -4) | ≥ 9 bp (-12 to -4) | 14 bp (-12 to +2) | 13 bp (-11 to +2) |
| W-dyad | | edge-on | edge-on | edge-on | edge-on | edge-on | chair |

### Supplemental Videos

**Supplementary Video 1.** Slow motion videos (240 fps; iPhone 8S) of the grid robot in the Spotiton device after initiation of plunges at the motion settings shown. The total plunge time is the entire time the robot is in motion. The reported mixing times begin when the grid has passed the second (lower) dispenser, visible in the lower part of the video, and ends when it enters the liquid ethane (not seen). Plunges shown are for demonstration only. No grid is held in the tweezer.

**Supplementary Video 2.** 3DVA video [1] showing the RNAP clamp open/close mode (dataset $4_{500ms}$). At the beginning of the video, the RNAP clamp is open. The clamp (light green labels) and $\sigma^{70}_{1.1}$ in the RNAP cleft (orange labels) are identified. As the clamp closes, $\sigma^{70}_{1.1}$ disappears and density corresponding to downstream duplex DNA appears (yellow labels). The video continues through three additional cycles of opening/closing.
